# Deep Learning on Chaos Game Representation for Proteins

**DOI:** 10.1101/575324

**Authors:** Hannah F. Löchel, Dominic Eger, Theodor Sperlea, Dominik Heider

## Abstract

Classification of protein sequences is one big task in bioinformatics and has many applications. Different machine learning methods exist and are applied on these problems, such as support vector machines (SVM), random forests (RF), and neural networks (NN). All of these methods have in common that protein sequences have to be made machine-readable and comparable in the first step, for which different encodings exist. These encodings are typically based on physical or chemical properties of the sequence. However, due to the outstanding performance of deep neural networks (DNN) on image recognition, we used frequency matrix chaos game representation (FCGR) for encoding of protein sequences into images. In this study, we compare the performance of SVMs, RFs, and DNNs, trained on FCGR encoded protein sequences. While the original chaos game representation (CGR) has been used mainly for genome sequence encoding and classification, we modified it to work also for protein sequences, resulting in n-flakes representation, an image with several icosagons.

We could show that all applied machine learning techniques (RF, SVM, and DNN) show promising results compared to the state-of-the-art methods on our benchmark datasets, with DNNs outperforming the other methods and that FCGR is a promising new encoding method for protein sequences.

## 1 Introduction

PProtein classification is one big challenge in bioinformatics [Heider et al., 2009], and has therefore many applications, ranging from genomic annotations towards clinical applications, such as drug resistance prediction in human immunodeficiency virus (HIV) for personalized therapies. To this end, different machine learning methods exist and have been applied, e.g., support vector machines (SVM) [Beerenwinkel et al., 2003], random forests (RF) [Heider et al., 2011, Löchel et al., 2018], or neural networks (NN) [Wang and Larder, 2003]. Generally, the protein sequences have to be made “machine-readable” in a first step. Different protein encodings exist, which can be roughly separated into sequence-based or structure-based encodings. These sequence-based encodings include sparse encoding [Hirst and Sternberg, 1992], amino acid composition [Matsuda et al., 2005], reduced amino acid alphabets [Solis and Rackovsky, 2000], physicochemical properties [Heider and Hoffmann, 2011], or Fourier Transformation [Nagarajan et al., 2006]. Structure-based encodings include quantitative structure-activity relationship (QSAR) [Cherkasov et al., 2014], Electrostatic Hull [Dybowski et al., 2011], or Delaunay triangulation Yu et al., 2013]. For a comprehensive review on encodings of protein sequences see Spänig and Heider [2019]. After encoding, the encoded sequences can be used for training of different machine learning models, such as SVMs, RFs, or deep neural networks (DNNs). Due to the fact that DNNs have been shown to outperform other methods in image classification, we will introduce a modified chaos game representation (CGR) for proteins and will show the performance of this encoding on HIV drug resistance prediction datasets in comparison to the state-of-the-art models. Moreover, we made our new frequency matrix chaos game representation (FCGR) for protein-encoding available as an R package kaos.

The chaos game representation (CGR) algorithm is a recurrent iterative function system, which can be used to create fractals from sequences of symbols, i.e., from an alphabet A={s1,…, sn}. For n=3 and A= {1,2,3}, the CGR algorithm can be used to construct, e.g,. the Sirpinski triangle, a fractal structure constructed by smaller triangles [Barnsley, 2012]. Jeffrey [1990] was the first who applied the CGR algorithm to DNA sequences, i.e., n=4 and A={A,C,G,T}, thus the resulting fractals are constructed from squares instead of triangles. The underlying idea of the CGR algorithm for DNA is summarized in Figure 1. Each symbol is set at one corner (here: 4). Starting from the middle, the next dot is put half the way towards the next symbol in the sequence. The second (and all remaining dots) are put half the way from the last position in the direction to the next symbol (exemplarily shown in Figure 1B.

**Figure 1:**
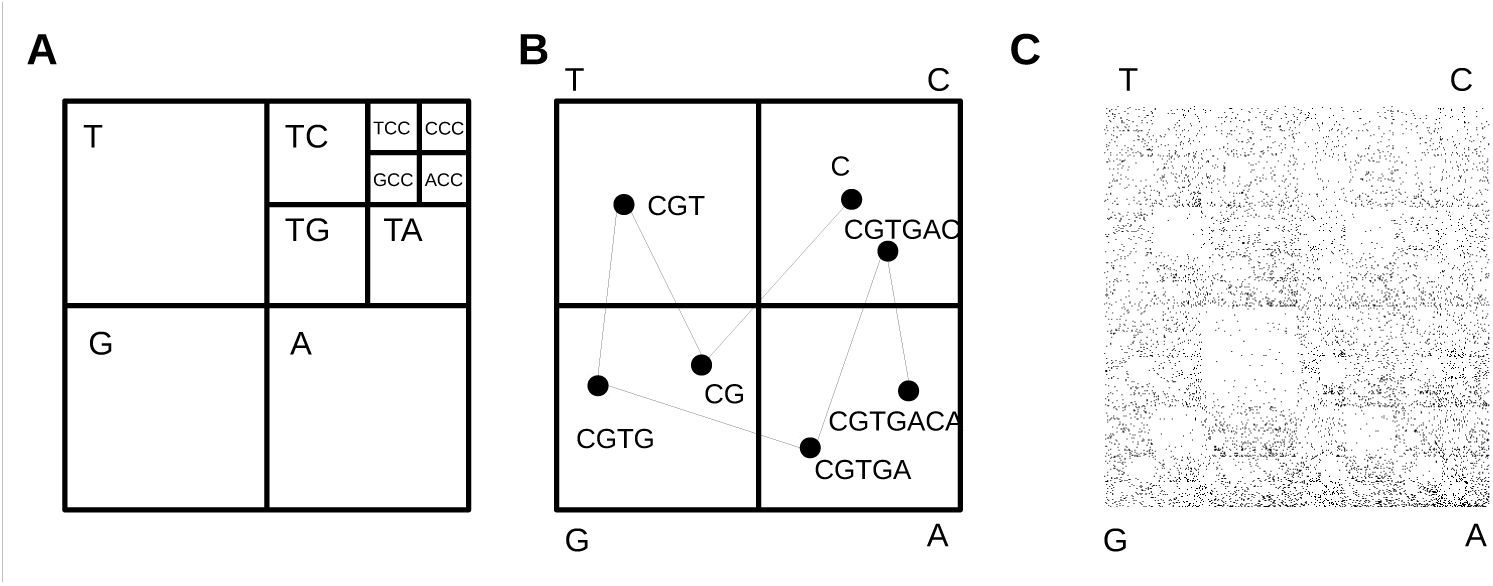
Chaos Game Representation for DNA. A) Division of the square. B) Way walked to draw points. C) HIV genome (NCBI Reference Sequence: NC 001802.1)

Since the development of the CGR and its application in life science, it has been used mainly for the analyses and comparison of whole genome sequences [Joseph and Sasikumar, 2006]. It has been shown that CGR is an excellent representation for genomes and that CGR-driven phylogeny leads to reliable predictions [Deschavanne et al., 1999]. In particular the comparison between genomes by using CGR is very easy and fast [Hoang et al., 2016]. Extensions of CGR include color grids [Deschavanne et al., 1999] and frequency matrix CGR (FCGR) [Almeida et al., 2001]. Wang et al. [2005] used FCGR to calculate the image distance between genomes in order to generate phylogenetic trees. Rizzo et al. [2016] showed that DNNs trained on genomes encoded with FCGR yielded very accurate predictions. They used a convolutional neural network (CNN) to divide bacteria in three different phyla, order, family, and genus and showed a very high accuracy for the method. While these studies focused only on FCGR for DNA, there exist also a smaller number of studies dealing with the encoding of protein sequences. Yu et al. [2004] employed the CGR algorithm for protein classification by separating the amino acids in four groups based on their properties and used multifractal and correlation analysis to construct a phylogenetic tree of Archaea and Eubacteria. In another approach the amino acids were retranslated into DNA for CGR [Yang et al., 2009]. Basu et al. [1997] used CGR by grouping the amino acids in twelve groups and used a twelve-sided regular polygon for the representation. Most of the studies with CGR on proteins have in common that they make use of the original approach to create the CGR, i.e., they go half the way of the distance to the next symbol to produce the CGR images. However, by using this approach, resulting CGR images are very noisy for alphabets with *n >* 4. In this study, we introduce the use of Sierpinskin-gons, also known as, n-flakes or polyflakes [Tzanov, 2015], which can be constructed by varying the distances and thus result in well-structured fractals. Moreover, we will make use of DNNs and FCGR for proteins and analyze the impact of the scaling factor as well as the resolution on the classification performance on HIV drug resistance datasets.

## 2 Methods

### 2.1 Dataset

HIV-1 is known for its high mutation rate, which offers the virus the opportunity to quickly evolve drug resistance. Thus, prediction of drug resistance is crucial for personalized therapy of the patient. Protein sequences of the HIV-1 protease (PR) and reverse transcriptase (RT) originating from subtype B strains with data for seven PIs (RTV: Ritonavir, IDV: Indinavir, SQV: Saquinavir, NFV: Nelfinavir, APV: Amprenavir, ATV: Atazanavir, LPV: Lopinavir), three NNRTIs (NVP: Nevirapine, EFV: Efavirenz, DLV: Delavirdine), and five NRTIs (3TC: Lamivudine, ABC: Abacavir, AZT: Zidovudine, D4T: Stavudine, DDI: Didanosine) with *IC*_50_ ratios were collected from the HIV Drug Resistance Database [Rhee et al., 2003]. The data was separated into susceptible and resistant by drug-specific cutoffs. Rhee et al. [2006] We removed sequences from the datasets for which no resistance information was available and excluded ATV from our classification approach, since too many sequences lacked *IC*_50_ information. Table 1 gives a summary of the data used in the study for each drug.

**Table 1:**
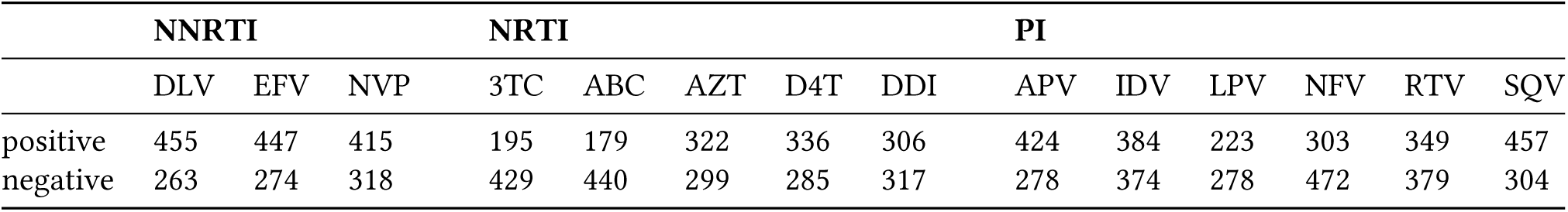
Data used in the study

### 2.2 Implementation of the chaos game representation algorithm

We implemented an R package kaos (downloadable from CRAN), which can be used to create CGR and FCGR with n-flakes. The kaos package accepts any kind of alphabets and creates the (F)CGR image based on the given sequence and user-specified resolution. The package offers the options to create an CGR image with dots (option “points”) or an FCGR (option “matrix”) with different gray-levels. For the FCGR, the user has to specify a resolution to specify the columns of the matrix. It is also possible to set the scaling factor (“sf”) which is needed to construct n-flakes. For protein sequences with twenty proteinogenic amino acids, the CGR representation results in twenty edges and twenty icosagons within a larger icosagon. The contraction ratio between the outer and the inner polygon can be calculated by the following equation [Strichartz, 2000]:

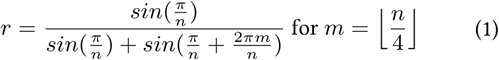

The ratio for the distance between the actual point and the target edge (i.e., the scale factor sf) can be calculated by the the following equation:

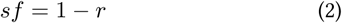

By default, the CGR package automatically creates the alphabet based on the given symbols or words in the sequence (vector of symbols or words) and takes this number as n to calculate the scaling factor by equation 1. The number is also needed to calculate the coordinates for the edges of a polygon in an unit circle with the following equation:

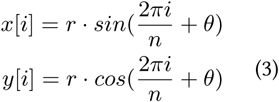

i: edge; n: number of edges, *θ*: angel of orientation

An CGR object contains the gray-level matrix with given resolution as an encoding for further analyses. In case of n=4, the CGR algorithm fills the whole matrix, otherwise it uses the unit circle.

Figure 2 shows examples created with the CGR package, namely the FCGR representation of the genomic DNA sequence of HIV with a resolution of 200, of the HIV RT sequence with a resolution of 50 and *sf* = 0.5, as well as of the HIV RT sequence with a resolution of 20 and *sf*_20_, the scaling factor for protein sequences with n-flakes. As mentioned before, the scaling factor is crucial in order to structure the fractal, which can be clearly seen by comparing the two FCGR representations. The CGR package offers predefined alphabets for numbers between 0-9, amino acids, DNA, and for the letters a-z as capital and lowercase letters.

**Figure 2:**
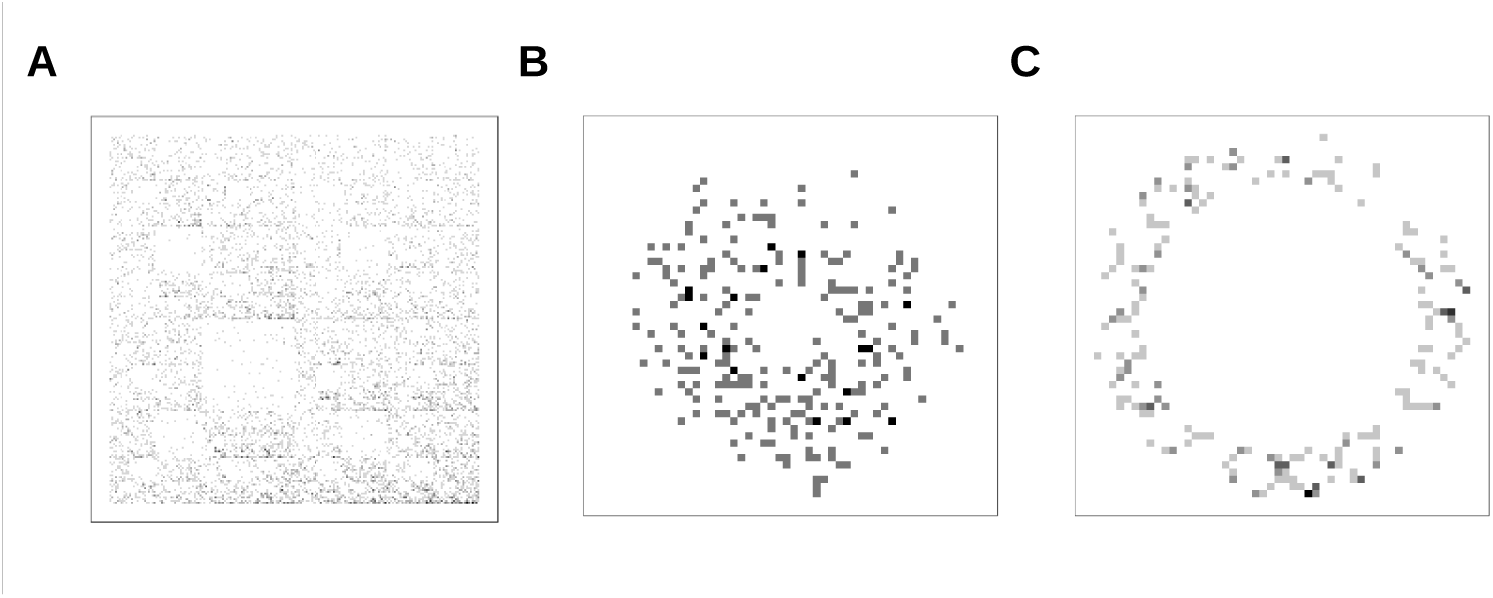
A) FCGR of genomic DNA sequence of HIV (NCBI Reference Sequence: NC 001802.1) with resolution of 200. B) AAQ18891.1 reverse transcriptase, partial [Human immunodeficiency virus 1] with resolution of 50 and *sf* = 0.5, C) AAQ18891.1 reverse transcriptase, partial [Human immunodeficiency virus 1] resolution = 50, *sf*_20_

### 2.3 Development of prediction models

In order to evaluate the impact of the resolution and the scaling factor on subsequent classification, we used eight different configurations for the CGR images and trained DNNs, RFs, and SVMs, with the settings for protein sequences (“amino”), to force 20-edges (see figure 3).

**Figure 3:**
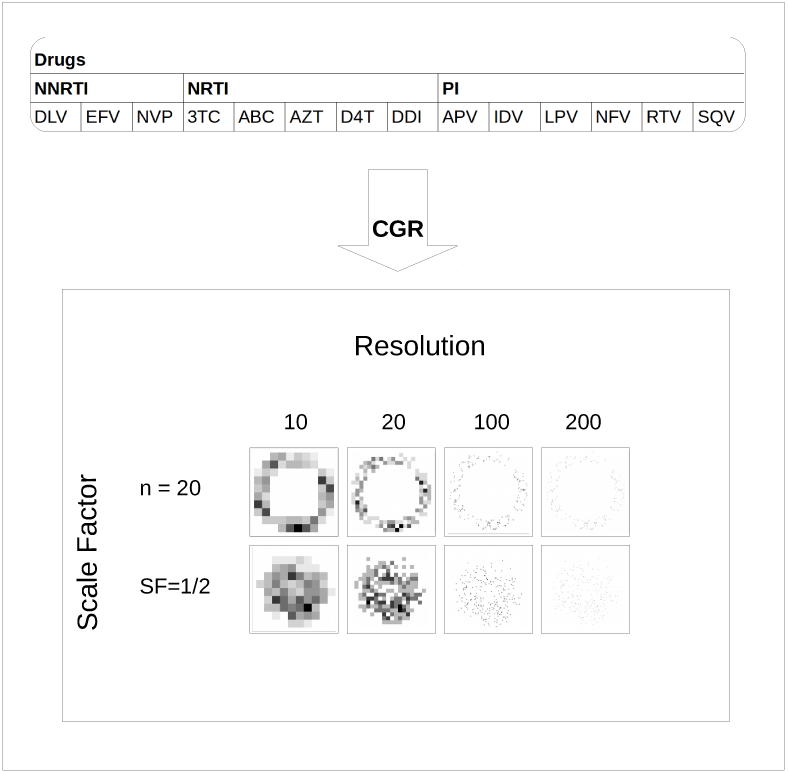
Different settings for producing the FCGR pictures. We used different combinations of resolution and scale factor to produce the FCGR images. The resolution was set to 10, 20, 100, and 200, while the scaling factor was set to 0.5 and *sf*_20_, i.e., the optimal scaling factor for n=20.

We performed a stratified hold-out validation scheme where 20 % of the data was randomly selected for validation and 60 % was used for building the models to evaluate the machine learning models. The remaining 20 % of the data was used as test data for the DNNs. We did not take this data for the SVMs and RFs, due to the fact that we wanted to keep the training data consistent with the DNNs. We then performed a 10-fold cross-validation with the remaining data (i.e., without the validation data). We trained models for SVMs, RFs, and DNNs with the different configurations mentioned before. All cells containing only zeros in all data were removed prior training of the SVMs and RFs.

For the SVMs we applied the e1071 package [Meyer et al.,2019] with the linear kernel and default settings, for the RFs the randomforest package [Breiman, 2001] with default settings and 1000 trees, and for the DNNs the deepnet package [Rong, 2014] in R. We trained the DNNs with *tangens hyperbolicus* as activation function. In addition, we varied the numbers of neurons (up to 20) in three hidden layers and number of training epochs (see Figure 4).

**Figure 4:**
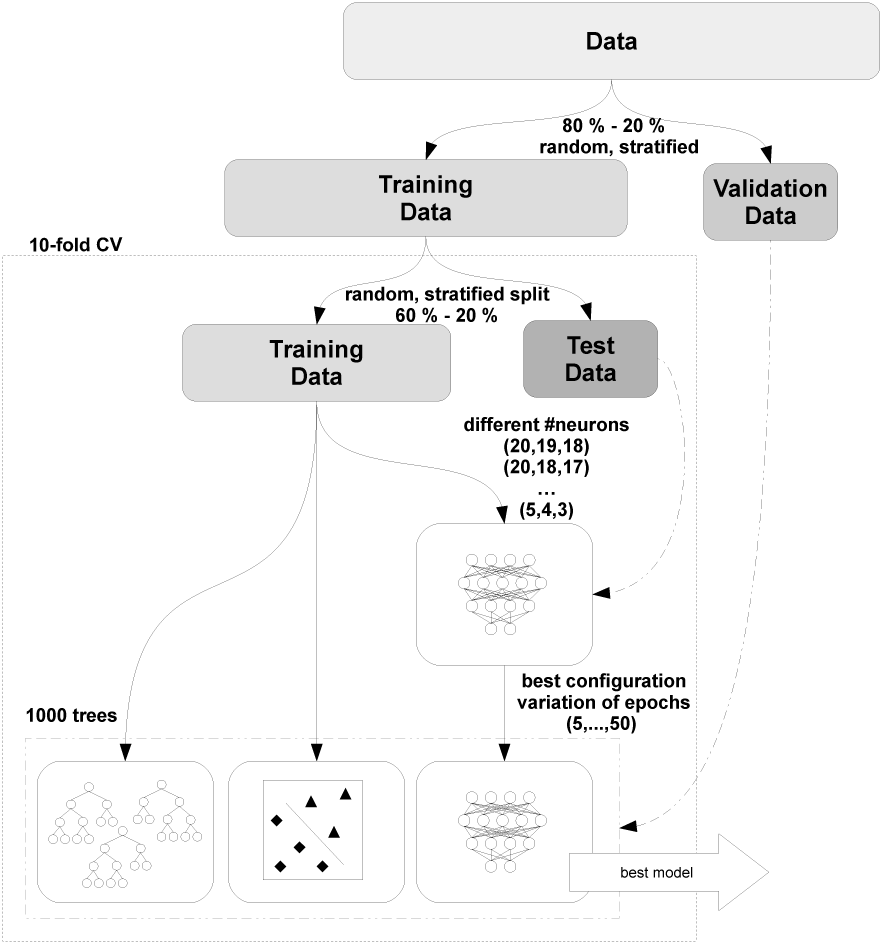
Development of prediction models.

We evaluated and compared the models based on the area under the receiving operating characteristics curve (AUC) with the R package pROC [Robin et al., 2011]. The best hidden layer configuration was selected based on the best average AUC. For the DNNs, we calculated the AUC also for the varying training epochs. Moreover, we used the R package ROCR [Sing et al., 2005] to draw precision-recall curves for the best-performing models.

### 2.4 Evaluation of FCGR as encoding

We calculated the average FCGRs of positive and negative samples, i.e., the average for each cell in the FCGR matrices of positive and negative sequences, respectively, in all datasets. Next, we calculated the differences between the average FCGR of the positive and the average FCGR of the negative samples. Significance of the differences were calculated based on Student’s t-tests, resulting p values were corrected for multiple testing by the method of Bonferroni. Moreover, in order to visualize the predictive quality of the different encodings in a model-independent manner, we used *< ϕ, δ >* diagrams as implemented in the R package phiDelta [Armano and Giuliani, 2018]. For this purpose we plotted the *< ϕ, δ >* diagrams for the encoding used by Heider et al. [2011] and the FCGR encoding with the different settings used in the current study. *< ϕ, δ >* diagrams are two-tiered 2D tools, which have been devised to support the assessment of classifiers or features in terms of accuracy and bias.

## 3 Results

We calculated the AUCs for the DNNs with different number of neurons from the cross-validation. For the best performing DNNs, we also evaluated different number of training epochs. Final evaluation of the models was carried out using the validation set. The best DNN configuration (number of neurons and epochs) in comparison to SVMs and RFs for the NRTIs and NNRTI, and the PIs are shown in figure 5 and figure 6, respectively. For the NRTIs ABC, DDI, and 3TC the DNN outperforms the other method in all encoding configuration, i.e., independently from resolution of the FCGR. However, for the NNRTIs DLV, EFV, and NVP, as well as for the NRTIs AZT and D4T, linear SVMs give better prediction results for lower resolutions. For very low resolutions, i.e., 10 and 20, the models are better with a scaling factor of 0.5. In all cases, the accuracy of the DNN models increases with the resolution. Figure 6 shows the results for the PIs. For high resolutions, the DNNs outperform the RFs and the SVMs, comparable to the results of the NNRTIs and NRTIs. While the performance of the DNNs increases with resolution, SVMs and RFs exhibit the opposite behavior. For low resolutions DNNs also outperform SVMs with one exception for SQV, where some SVM models perform better at a resolution of 20 and a scaling factor for n = 20 than DNNs.

**Figure 5:**
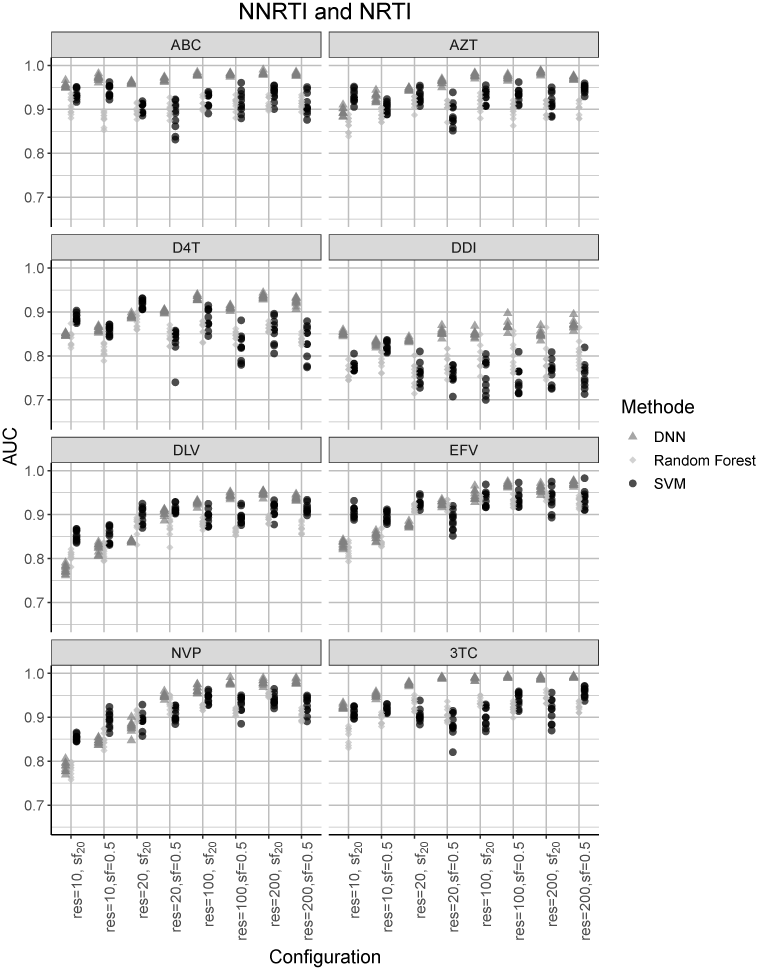
AUCs for NNRTIs and NRTIs. Results from with different training splits and different configurations evaluated with the validation data. Triangle: DNN; raute: RF; circle: SVM.

**Figure 6:**
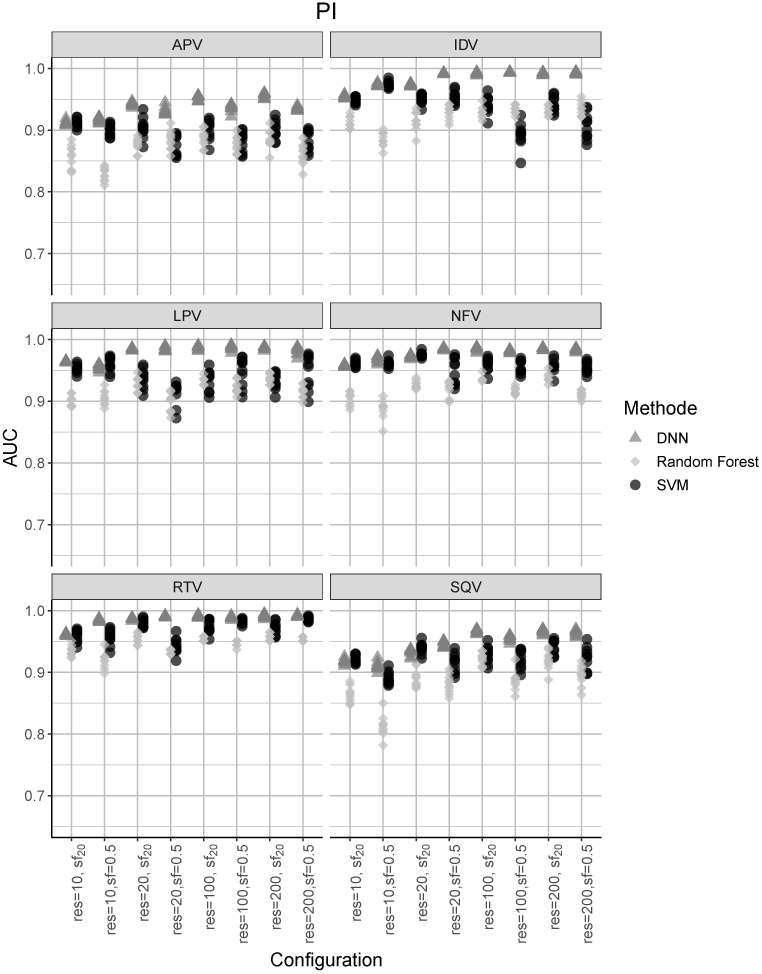
AUCs for PIs. Results from with different training splits and different configurations valuated with the validation data. Triangle: DNN; raute: RF; circle: SVM.

In Table 2 the AUCs of the best RF, SVM, and DNN models are summarized. The DNNs outperform all other methods, except for EFV, where the SVM performs equally well at a resolution of 200. For the PIs, best results are also observed with the DNNs, in the most cases with *sf*_20_, except for IDV and LPV, where the best result is observed at a scaling factor of 0.5. For high resolution the DNNs work best. The optimal scaling factor depends on the dataset, e.g., for APV there are higher AUC values with *sf*_20_, however, for DDI the best results are obtained with *sf* = 0.5. Some datasets perform quite well at low resolution, especially ABC and RTV, whereas increasing the scaling factor has a barely remarkable influence on the AUC values. While the DNNs have the highest AUC values, the other models still perform quite well, and thus supports the idea of CGR for protein encoding. Figure 7 and Figure 8 show the precision-recall curves for the best DNNs for the different drugs, supporting the very good prediction results from the DNNs.

**Table 2:** Best AUCs for NNRTIs and NRTIs on validation data

|  | DNN | RF | SVM |
| --- | --- | --- | --- |
| **NNRTI** |  |  |  |
| DLV | 0.95 ($r=200$, $sf=0.86$) | 0.90 ($r=200$, $sf=0.86$) | 0.93 ($r=200$, $sf=0.5$) |
| EFV | 0.98 ($r=200$, $sf=0.5$) | 0.95 ($r=200$, $sf=0.86$) | 0.98 ($r=200$, $sd=0.5$) |
| NVP | 0.99 ($r=100$, $sf=0.5$) | 0.96 ($r=200$, $sf=0.86$) | 0.96 ($r=200$, $sf=0.86$) |
| **NRTI** |  |  |  |
| 3TC | 0.99 ($r=100$, $sf=0.5$) | 0.96 ($r=200$, $sf=0.86$) | 0.97 ($r=200$, $sf=0.5$) |
| ABC | 0.99 ($r=100$, $sf=0.5$) | 0.94 ($r=10$, $sf=0.86$) | 0.96 ($r=10$, $sf=0.5$) |
| AZT | 0.99 ($r=200$, $sf=0.86$) | 0.94 ($r=100$, $sf=0.86$) | 0.96 ($r=100$, $sf=0.5$) |
| D4T | 0.94 ($r=200$, $sf=0.86$) | 0.90 ($r=20$, $sf=0.86$) | 0.93 ($r=20$, $sf=0.86$) |
| DDI | 0.90 ($r=100$, $sf=0.5$) | 0.87 ($r=200$, $sf=0.86$) | 0.84 ($r=10$, $sf=0.5$) |
| **PI** |  |  |  |
| APV | 0.96 (200 $sf=0.86$) | 0.91 (200 $sf=0.86$) | 0.93 (20 $sf=0.86$) |
| IDV | 0.995 (200 $sf=0.5$) | 0.95 (200 $sf=0.86$) | 0.98 (10 $sf=0.5$) |
| LPV | 0.99 (100 $sf=0.5$) | 0.95 (200 $sf=0.86$) | 0.98 (200 $sf=0.5$) |
| NFV | 0.99 (100 $sf=0.86$) | 0.95 (200 $sf=0.86$) | 0.98 (20 $sf=0.86$) |
| RTV | 0.995 (200 $sf=0.86$) | 0.96 (200 $sf=0.86$) | 0.99 (200 $sf=0.5$) |
| SQV | 0.97 (200 $sf=0.86$) | 0.94 (200 $sf=0.86$) | 0.96 (20 $sf=0.86$) |

r: resolution; sf: scaling factor.

**Figure 7:**
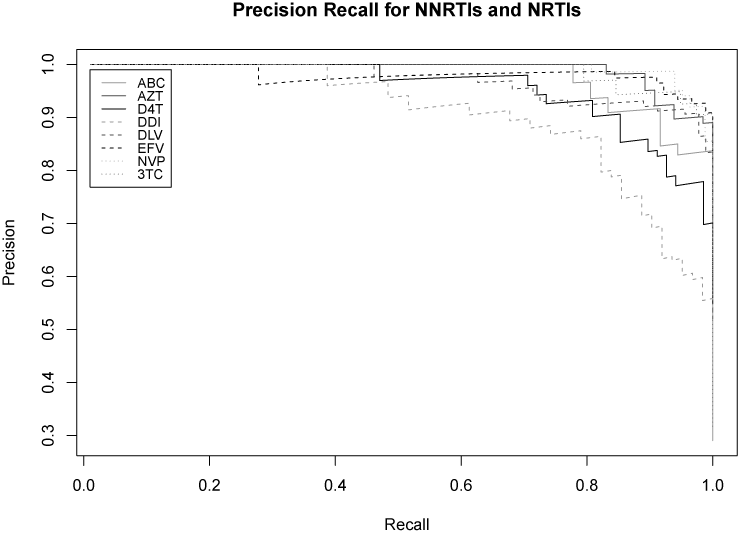
Precision-recall curves for PIs.

**Figure 8:**
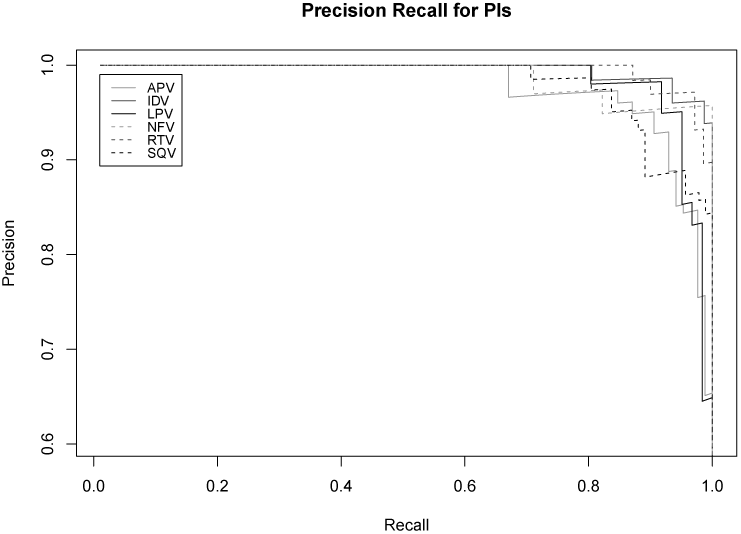
Precision-recall curves.

### 3.1 Comparison with other encodings

So far, we only compared the results from the different models, namely DNNs, SVMs, and RFs, on the same protein encoding, namely the FCGR. In the following, we will compare our results with the state-of-the-art methods.

Table 3 shows the AUC values of the best models trained on FCGR from our approach in comparison to the models of Heider et al. [2011] and Kierczak et al. [2009] for NRTIs and NNRTIs. Compared to the approach of Heider et al. [2011] and Kierczak et al. [2009], we get AUC values between 4 % up to 8 % and 19 % higher, respectively. Even the lower performing SVMs and RFs outperform or at least perform equally well compared the state-of-the-art approaches.

**Table 3:**
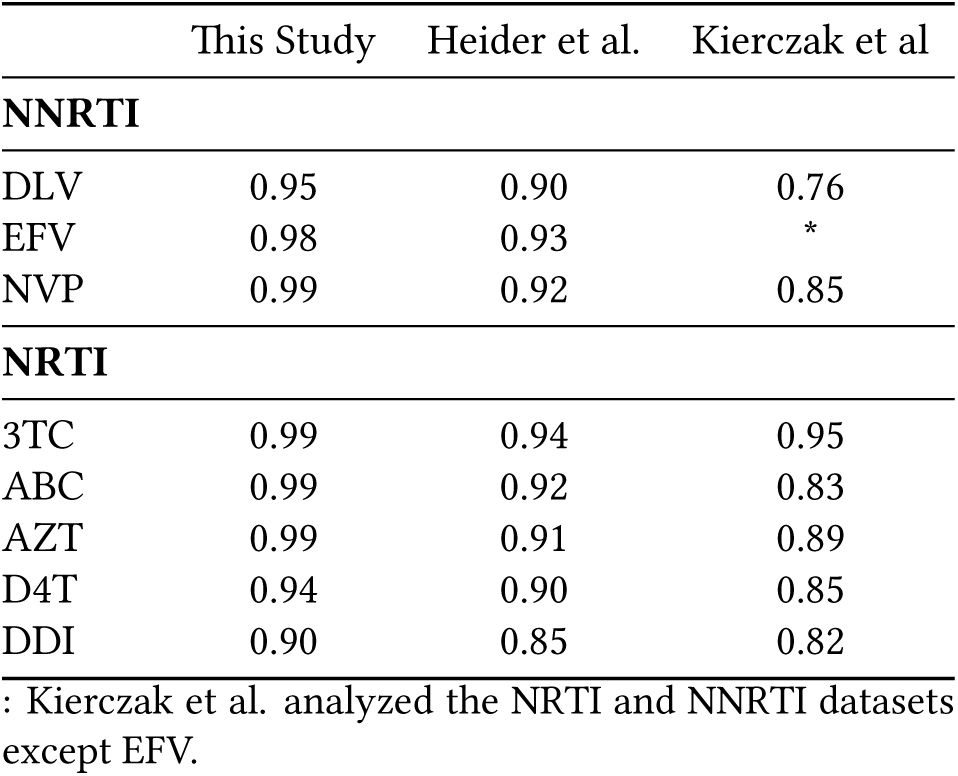
Comparison of the FCGR approach with state-of-the-art methods for NNRTIs and NRTIs

|  | This Study | Heider et al. | Kierczak et al |
| --- | --- | --- | --- |
| **NNRTI** |  |  |  |
| DLV | 0.95 | 0.90 | 0.76 |
| EFV | 0.98 | 0.93 | * |
| NVP | 0.99 | 0.92 | 0.85 |
| **NRTI** |  |  |  |
| 3TC | 0.99 | 0.94 | 0.95 |
| ABC | 0.99 | 0.92 | 0.83 |
| AZT | 0.99 | 0.91 | 0.89 |
| D4T | 0.94 | 0.90 | 0.85 |
| DDI | 0.90 | 0.85 | 0.82 |

: Kierczak et al. analyzed the NRTI and NNRTI datasets except EFV.

Table 4 shows the calculated accuracy values for the best models, in comparison with Heider et al. [2011], Rhee et al. [2006], and Hou et al. [2009] for the PIs. For all drugs, the FCGR approach outperforms the state-of-the-art models.

**Table 4:**
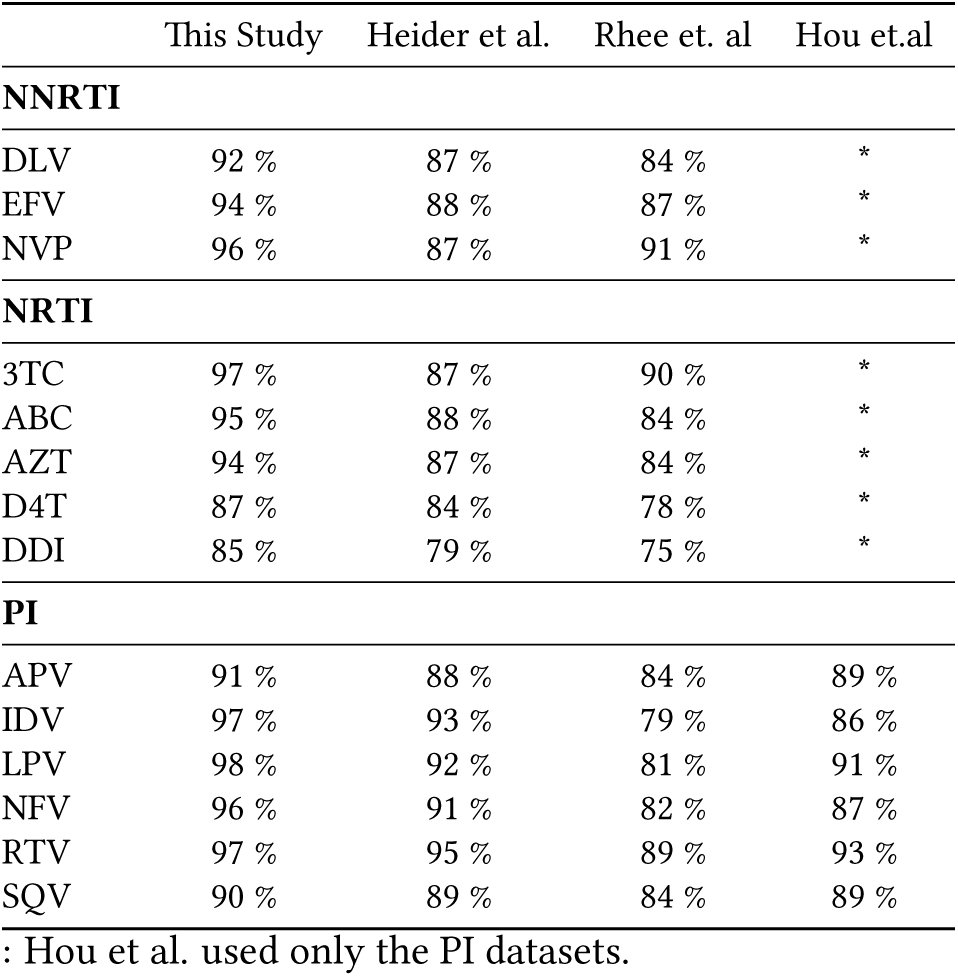
Comparison of the FCGR approach with state-of-the-art methods

The fact that FCGR-based classifiers were consistently out-performing other classification models in this study suggests that FCGR itself is a feature encoding for protein sequences preferable to some others. In order to test this hypothesis with regards to the data analyzed here, we compared the predictiveness of the feature encodings used in this study with the amino acid encoding and interpolation based feature encoding used in Heider et al., 2011 using *< ϕ, δ >* diagrams [Armano et al., 2018], which allow for the visual inspection of model-independent feature quality with regards to a given binary classification task (Figure 9 A and B). For all sequence datasets analyzed here, FCGR-based features show superior predictiveness (see supporting information). To explain this behaviour of FCGR encodings, we compared FCGR matrices for the positive and negative sequences from the different datasets. These show clear and significant differences in a small number of pixels (see supporting information). This is in accordance with the finding that very different machine learning models trained with FCGR-encoded sequences show consistently high performance.

**Figure 9:**
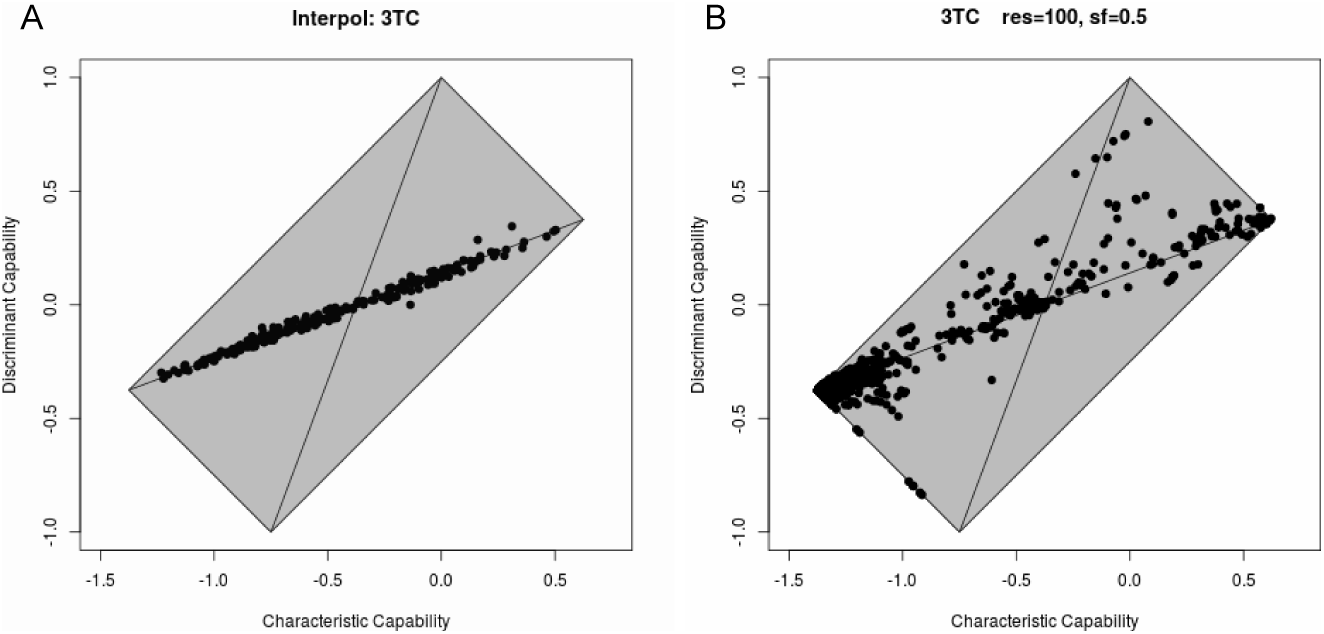
Comparison of amino acid and FCGR based feature representations. Exemplarily shown for the 3TC sequence dataset. A and B: *< ϕ, δ >* diagrams displaying the quality of features for the same sequence dataset (3TC) calculated using the R package Interpol Heider and Hoffmann [2011] as used in Heider et al. [2011] (A) and by FCGR with a resolution of 100 and *sf* = 0.5 (B). Dots represent features; dots closer to the upper or lower corner of the quadrilateral represent features with a high predictiveness for the classification task.

## 4 Discussion

The performance in terms of AUC of the RFs and SVMs has a higher variance compared to the AUCs of the DNNs, i.e., the split of test and training data might have a larger impact on the training of these models than on the DNNs. It can be observed that the DNNs perform better than the other models for higher resolution images. We can also observe that for some drugs low scaling factors work quite well and that the increase barely influences the results, whereas for other drugs the scaling factor leads to better performance until a saturation is reached. This suggests that the scaling factor somehow reveals patterns on some resolution, characteristic for the classification on this dataset. Comparing the course of the different models (Figure 5), we can see that the SVMs and DNNs perform equally good on a high level. Especially for the RFs we can observe that the application of the scaling factor increases the performance. There might be a saturation for the performance of the DNNs at a given resolution where the application of *sf*_20_ or using 0.5 has a low impact on the performance. We can observe this for most of the drugs. Except for D4T and DDI where there is a drop in prediction performance. The models trained on FCGR outperform all other evaluated models, independent of the employed machine learning technique. This suggests that FCGR as an encoding for protein sequences might be more appropriate than other encodings. By using the *< ϕ, δ >* diagrams we could show that the FCGR features show superior predictiveness. In comparison with the method of Heider et al. [2011], the FCGR encoding has no information loss on high resolution. Due to the interpolation the sequence-length is changed and this can lead to a loss of information. The advantage of the FCGR encoding is that the amino acid as itself is not transformed in any kind of representation, e.g., physicochemical properties. It can be considered as a kind of black box, where each letter represent different unknown feature lying behind each letter. The order of the letters is more or less kept, depending on the resolution, which explains the increase of performance in a higher resolution. One disadvantage is the increase of memory requirements for one FCGR matrix compared to a string or vector. In particular the use of *sf*_20_, where most of the space in an FCGR image is never used. Thus, a solution might be to finally erase those elements of the matrix. We used comparatively long protein sequences in this study, thus, one open question is, if the FCGR encoding still works well for shorter sequences, e.g., peptides, since the formation of patterns might be less pronounced for short sequences.

## 5 Conclusion

FCGR as a feature encoding for proteins reveals a new approach for classification problems, which is particularly well-suited for DNNs. The encoding shows superior behavior compared to other encodings, independent from the employed machine learning technique in our study dealing with HIV-1 drug resistance. In fact, it outperforms the state-of-the-art methods and therefore it might be preferable to other protein classification problems. In combination with DNNs, FCGR can give very accurate predictions. The application of the scaling factor, in order to make use of n-flakes for training, can increase the accuracy, especially for RFs. Besides, the resolution of the FCGR plays an important role and can increase the accuracy depending on the classification problem. Since the FCGR method offers the opportunity to encode all kind of sequences, e.g., text and numbers, the use of FCGR in many other kind of applications besides DNA and protein classification problems, might be reasonable.

## Supporting information

Phi-delta diagrams and CGRs for all drugs

