## Supplementary material for "Deep Learning on Chaos Game Representation for Proteins": Phi-delta diagrams and CGRs for all drugs

March 2019

**Abstract**

This sections contains the $< \phi, \delta >$ [Armano and Giuliani, 2018] the encoding used by Heider et al. [2011] for all drugs (Figure 1), and within the CGR encoding, for all configurations used in this study (left column Figure 3 to Figure 29). The difference between the average FCGR of positive and negative sequences in the datasets are shown in the middle column of Figure 3 to Figure 29, and the significance of the differences in log(p) values as calculated using a Bonferroni-corrected t-test in the right column. Blue pixels are significantly different between positively and negatively labeled sequences. Gray pixels contain no values or zeros.

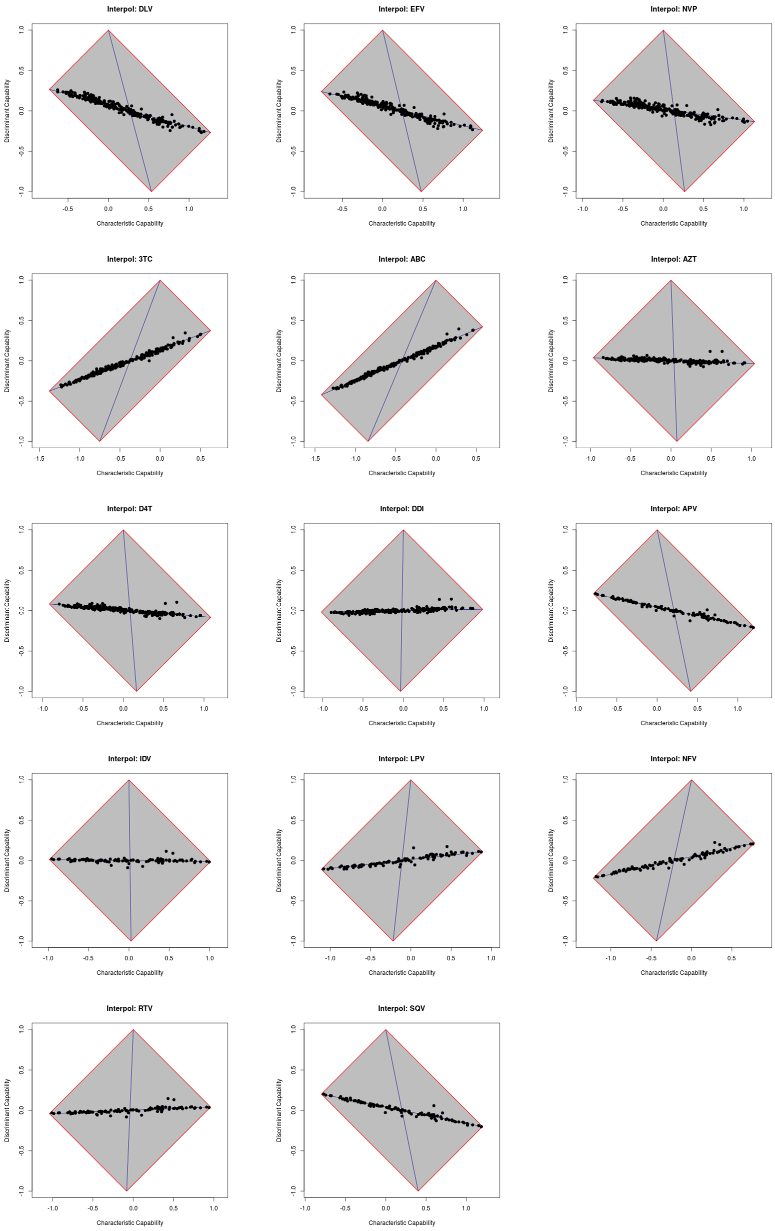


Figure 1: $< \phi, \delta >$ diagrams, calculated using the R package Interpol [Heider and Hoffmann, 2011]

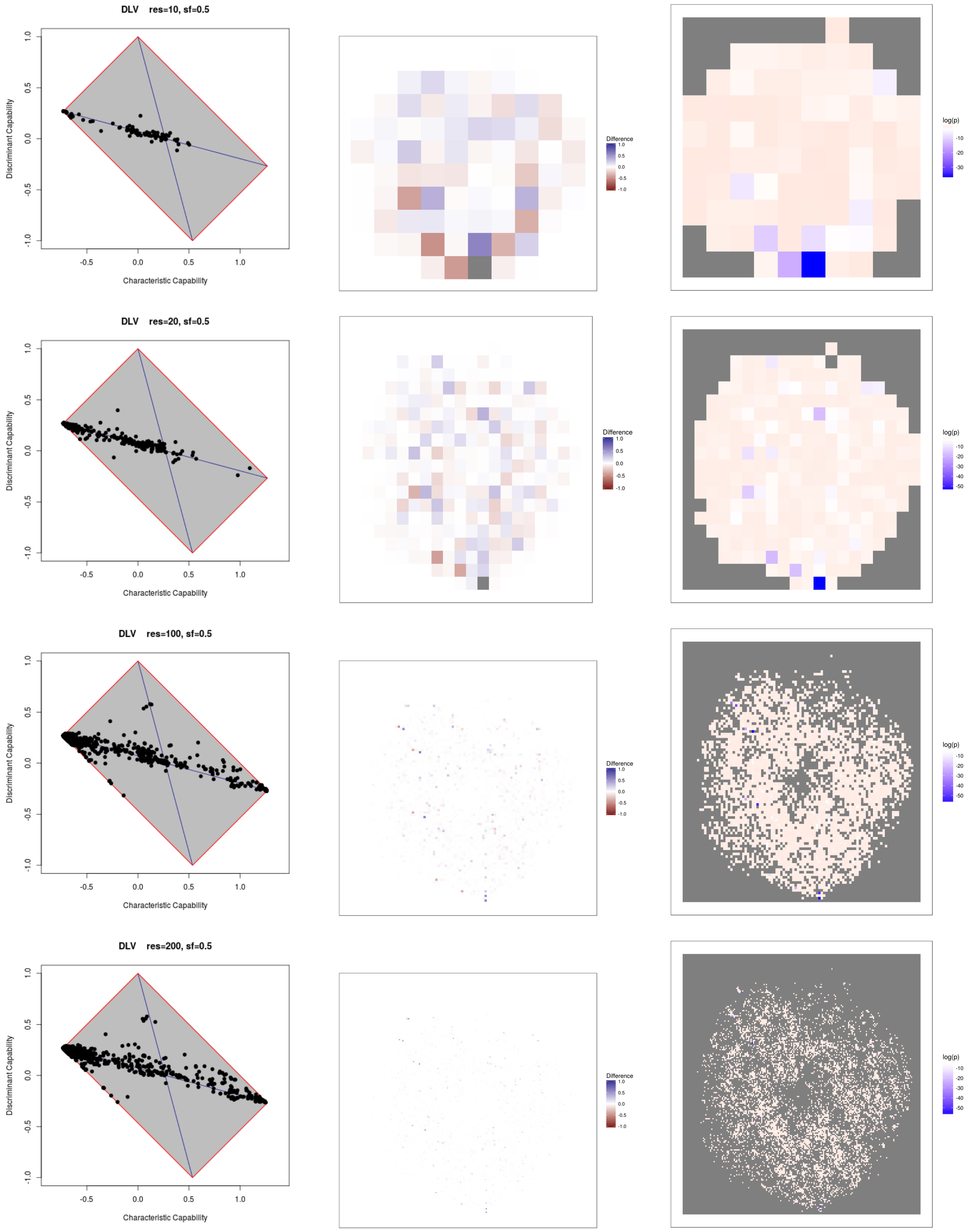


Figure 2: DLV, sf=0.5

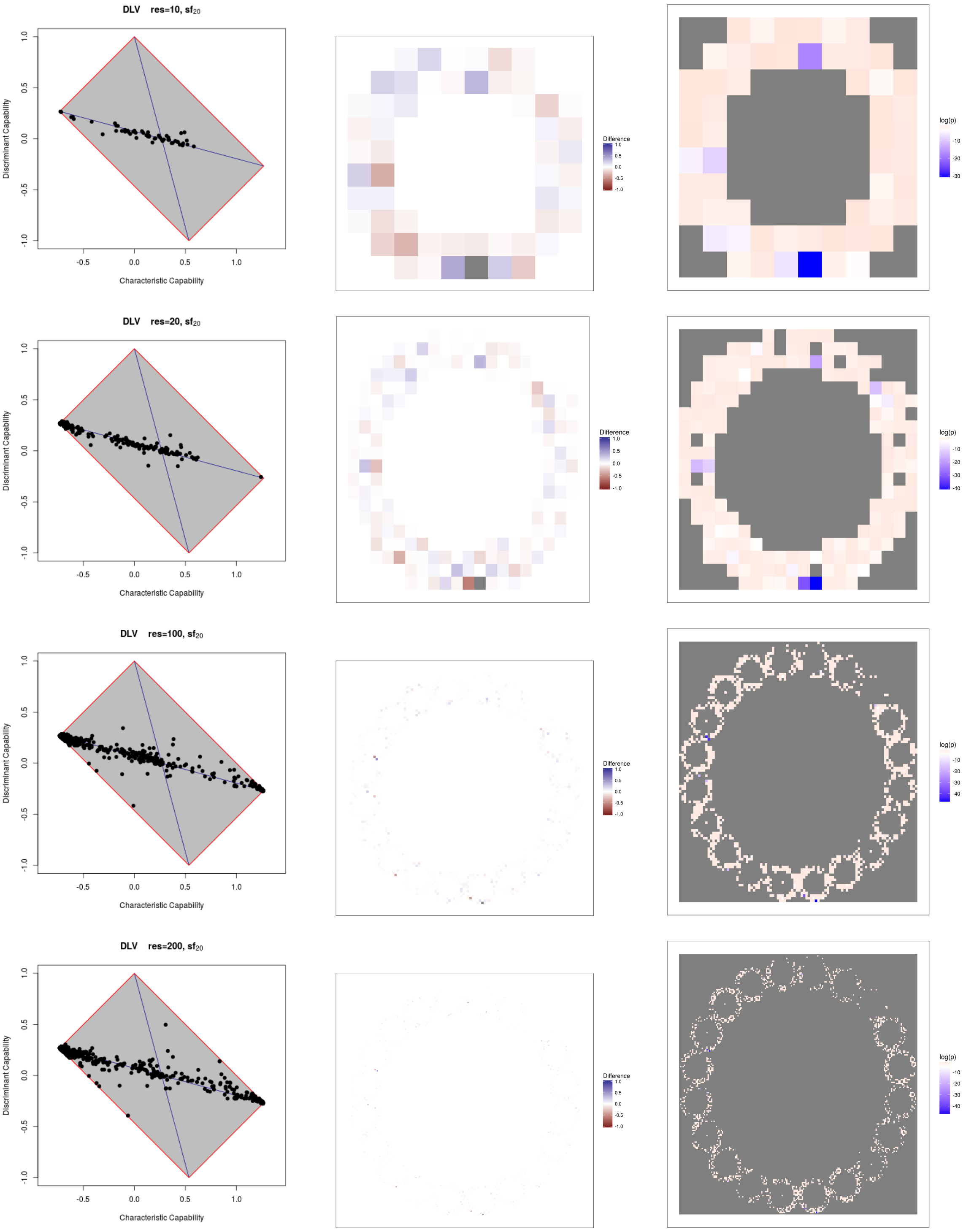


Figure 3: DLV, $sf_{20}$

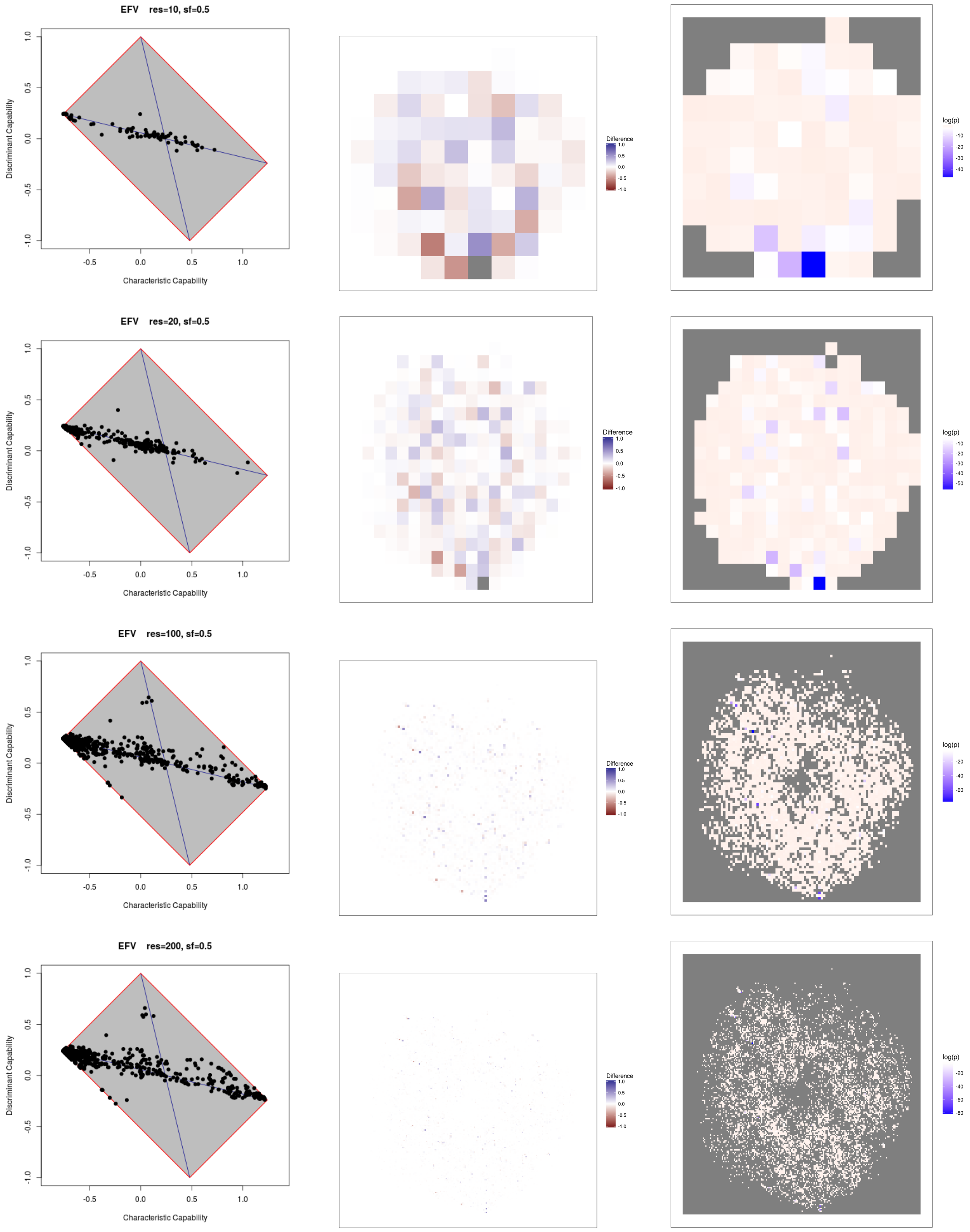


Figure 4: EFV, sf=0.5

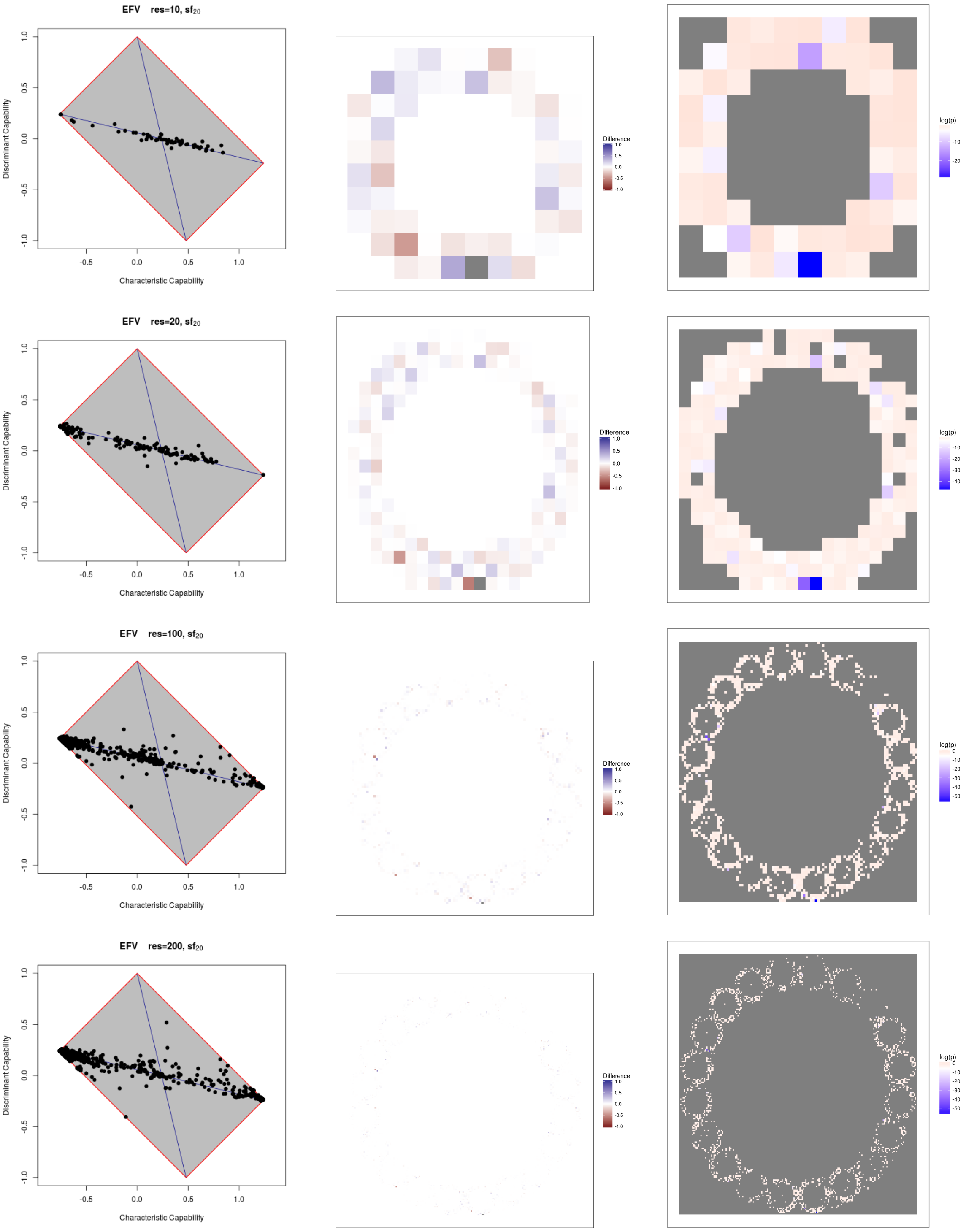


Figure 5: EFV, $sf_{20}$

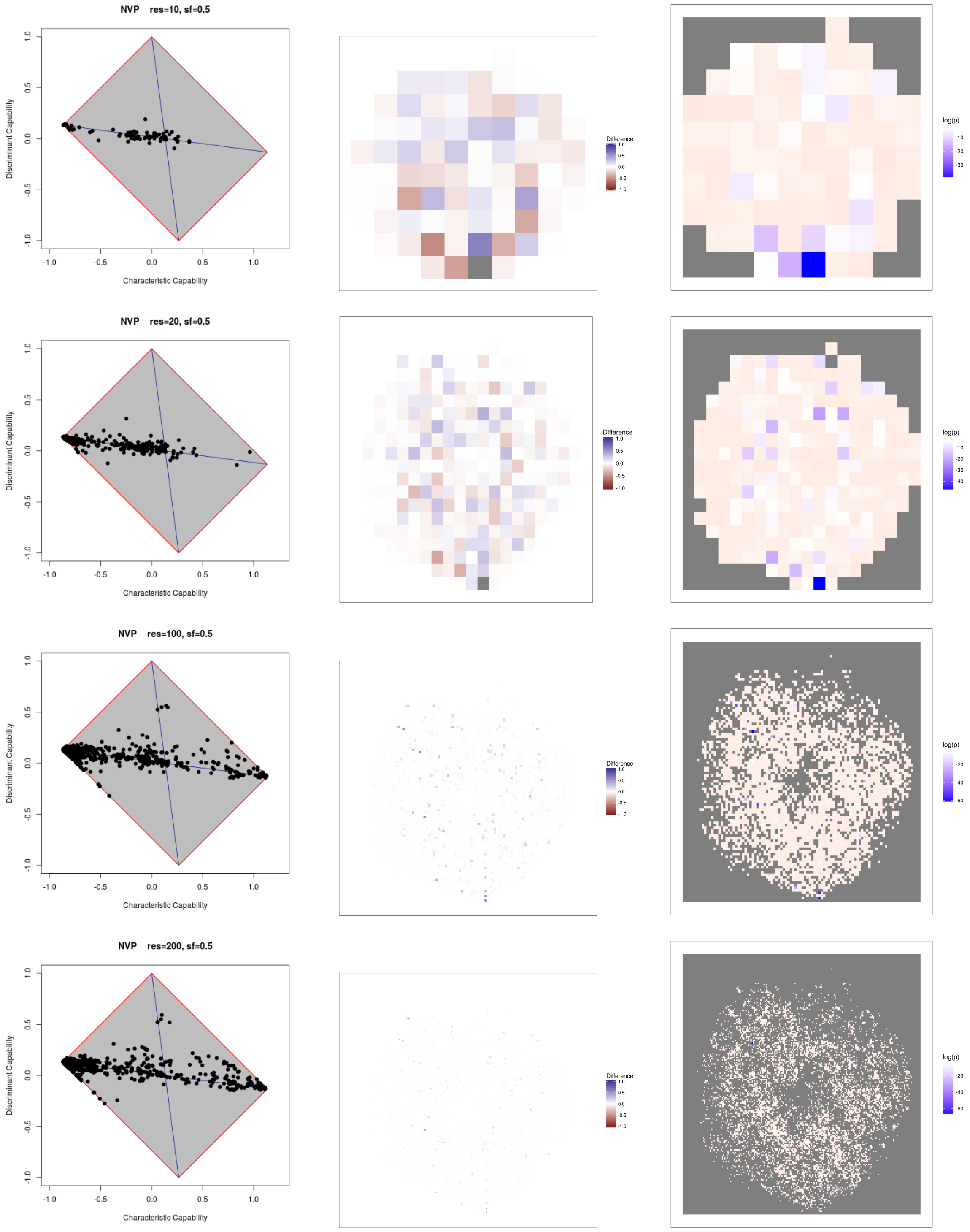


Figure 6: NVP, sf=0.5

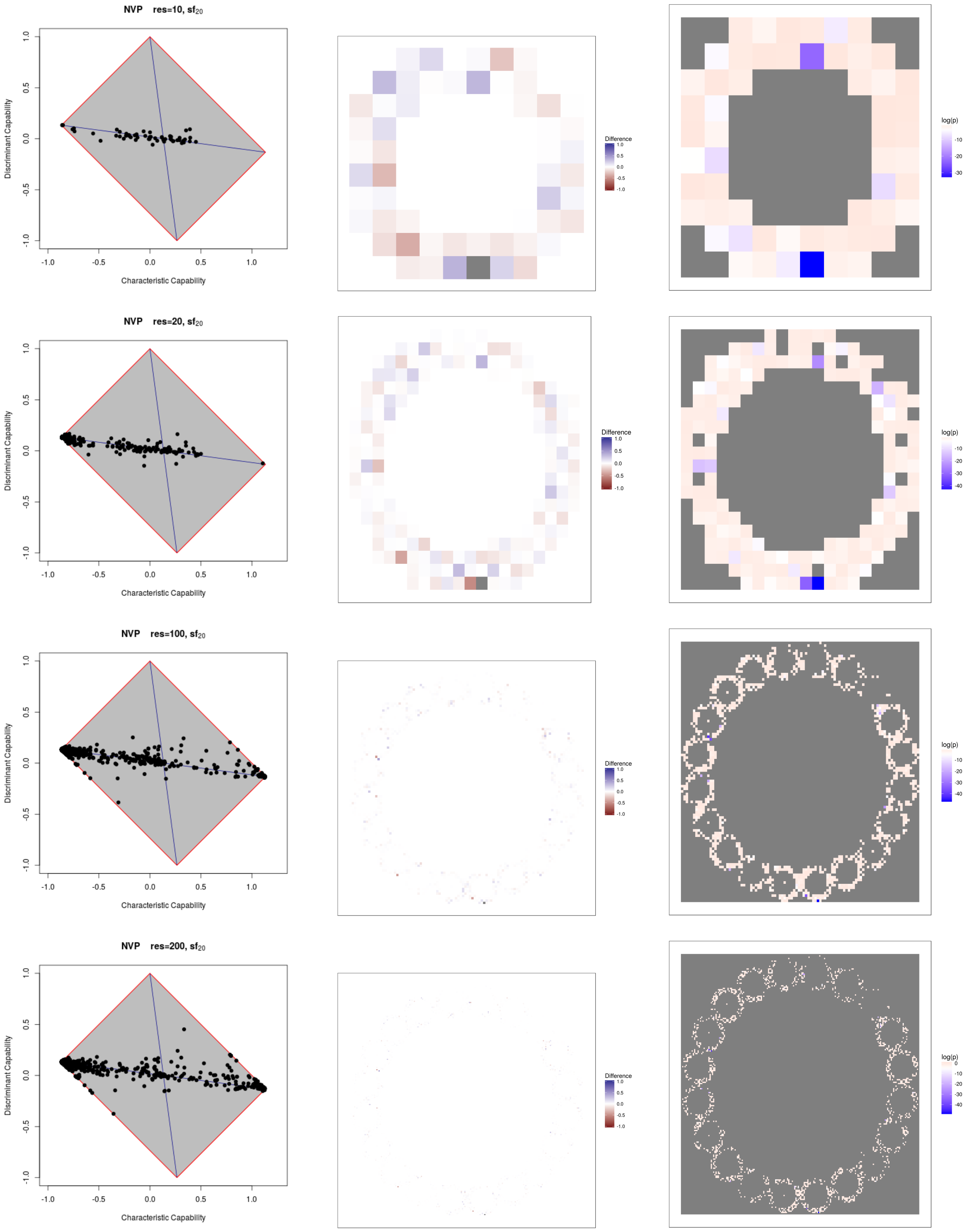


Figure 7: NFV, $sf_{20}$

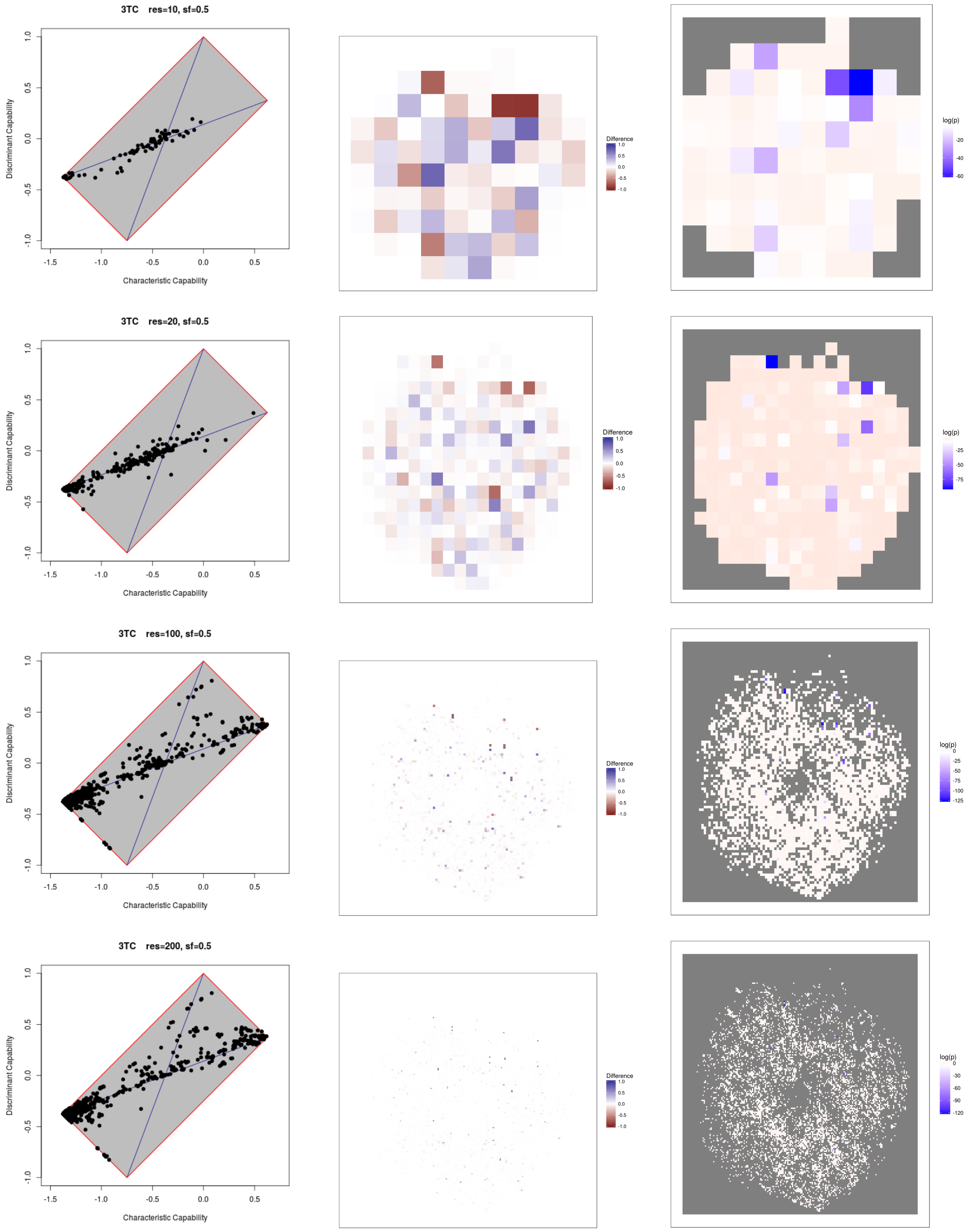


Figure 8: 3TC, sf=0.5

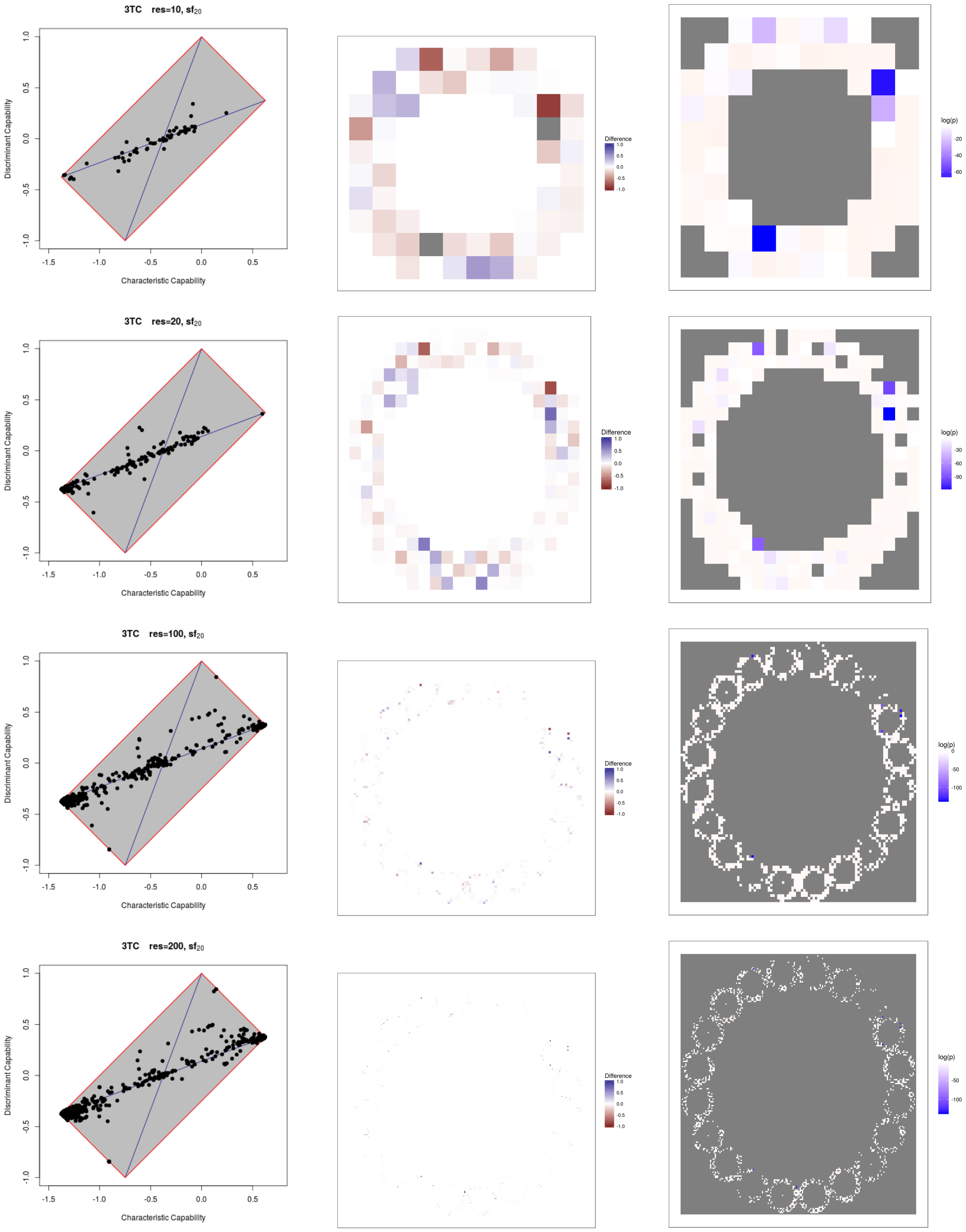


Figure 9: 3TC, $sf_{20}$

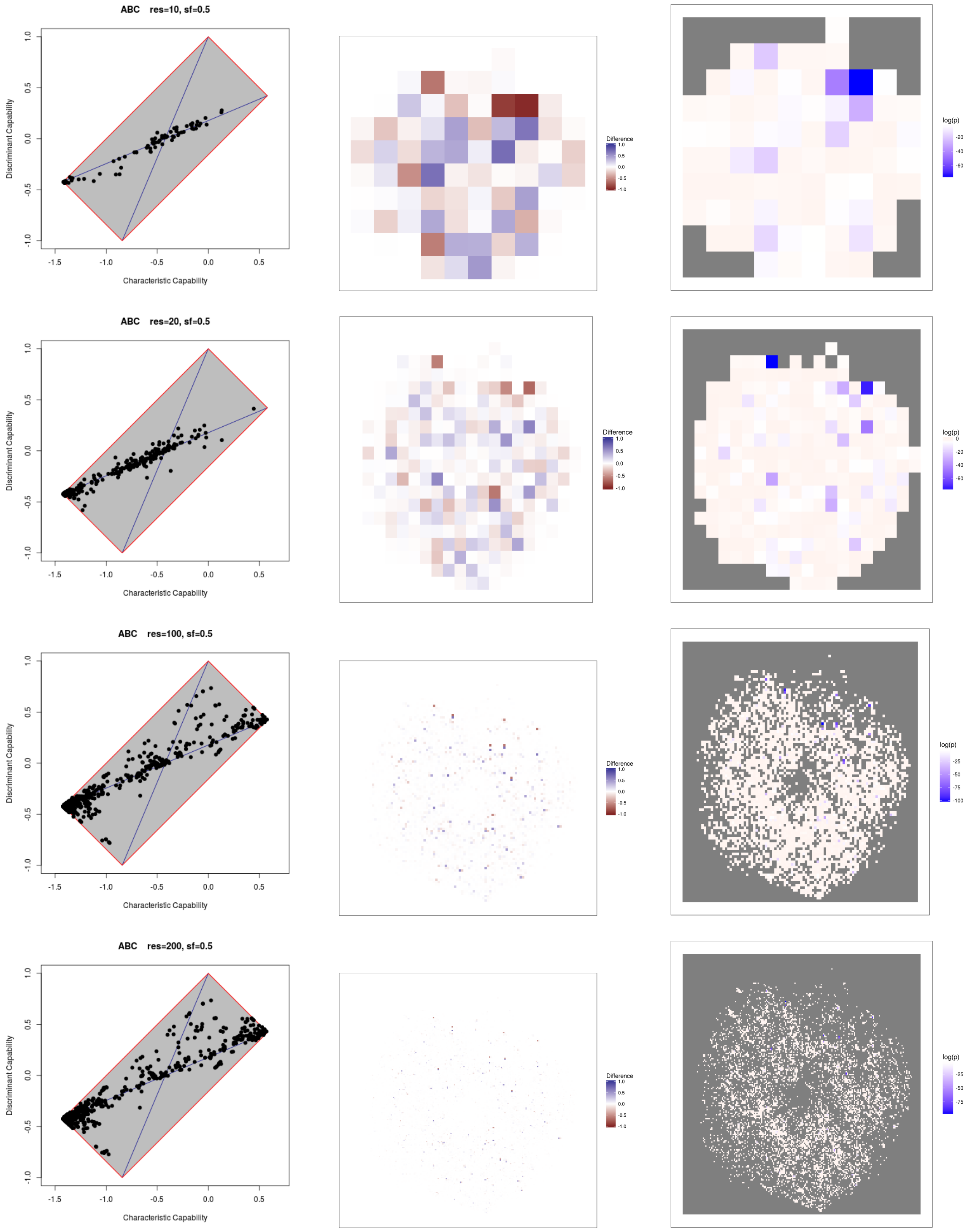


Figure 10: ABC, sf=0.5

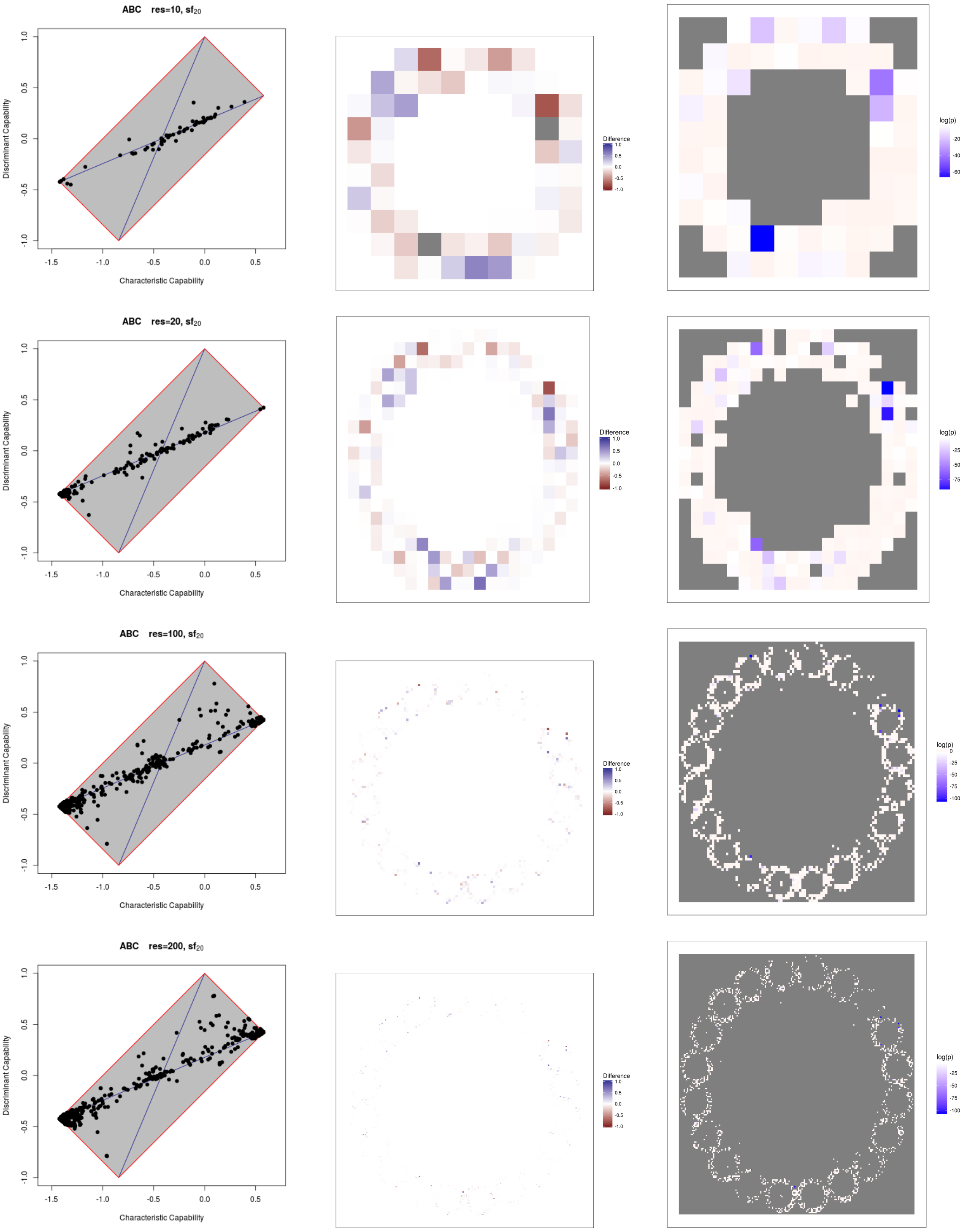


Figure 11: ABC, $sf_{20}$

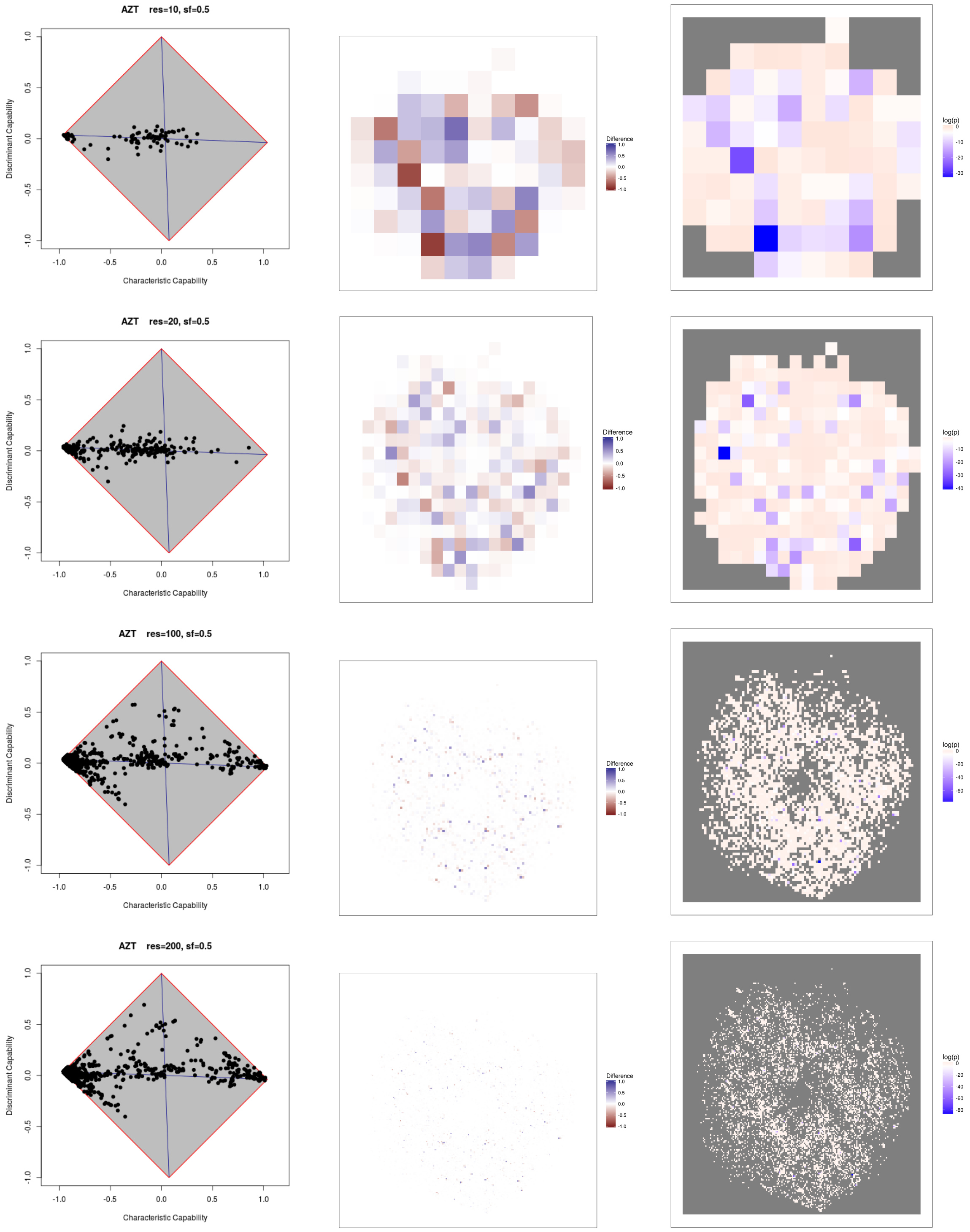


Figure 12: AZT, sf=0.5

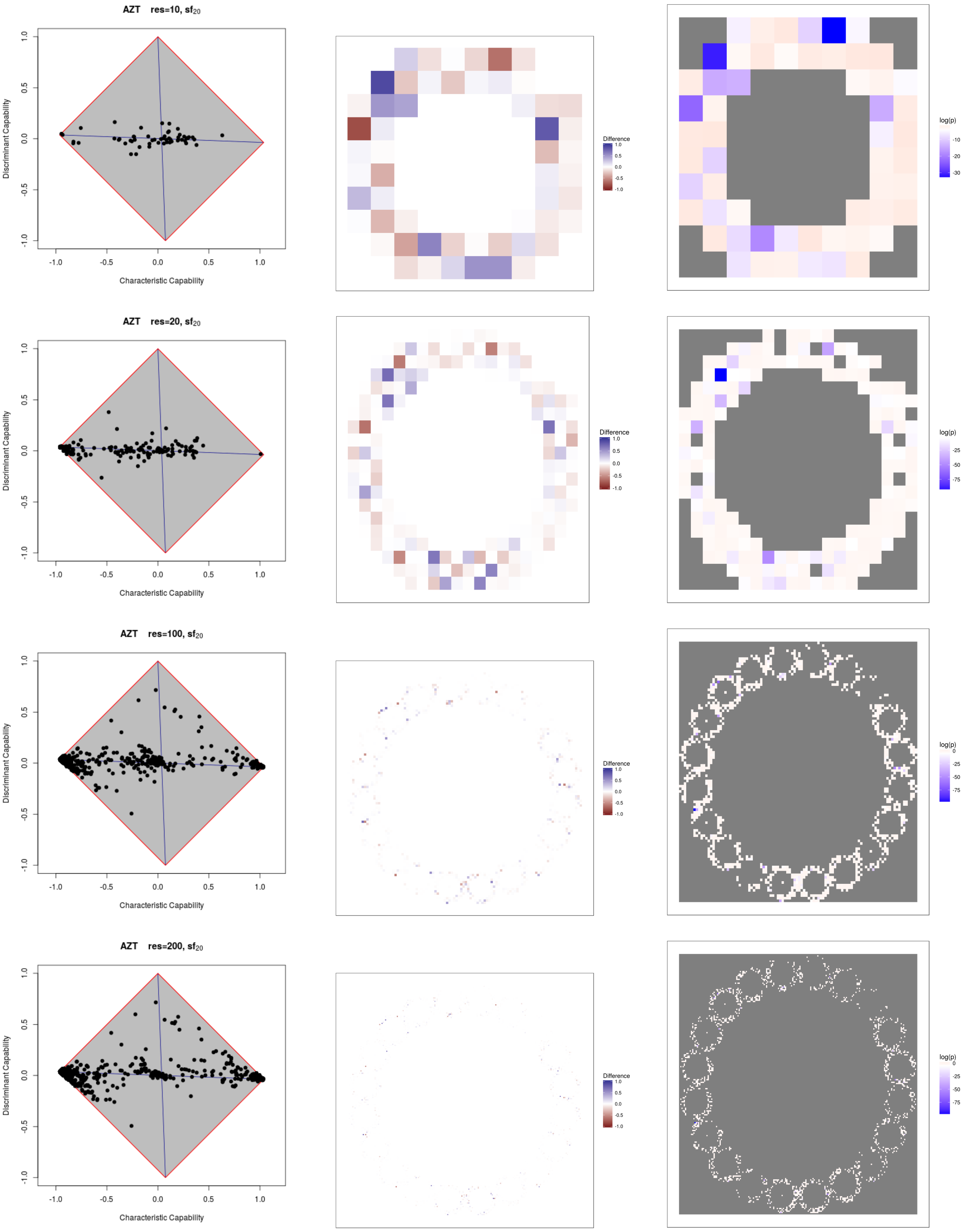


Figure 13: AZT, $sf_{20}$

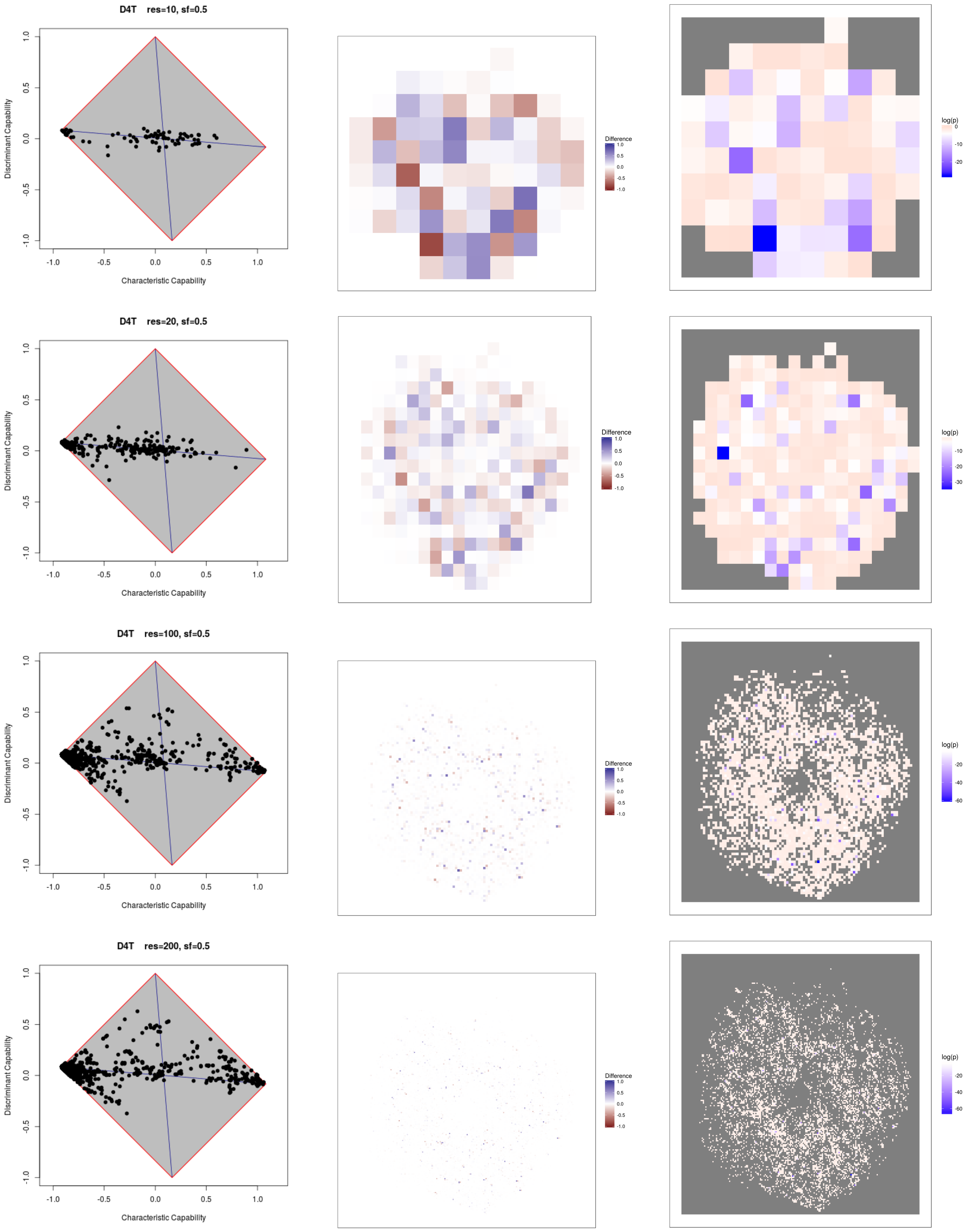


Figure 14: D4T, sf=0.5

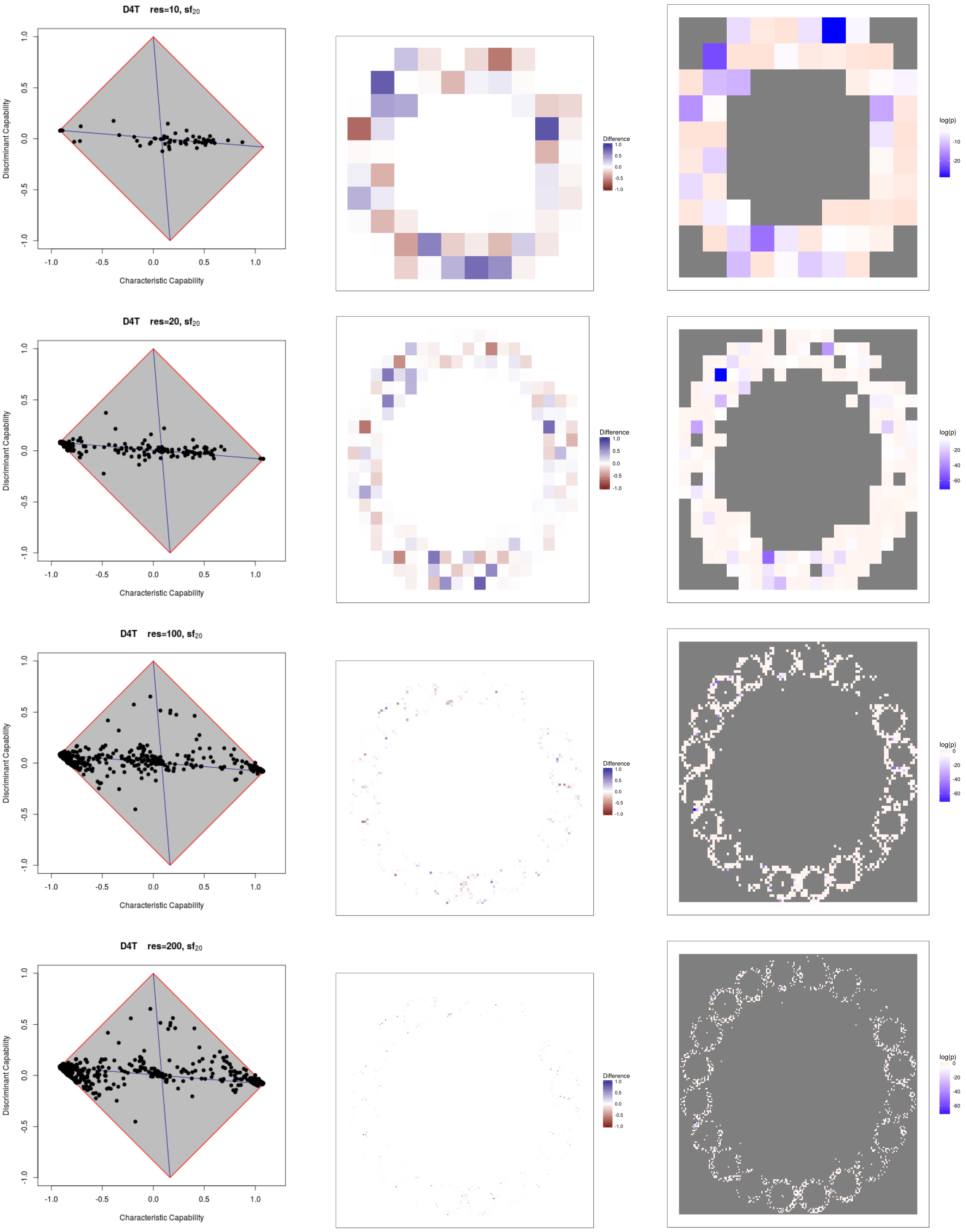


Figure 15: D4T, $sf_{20}$

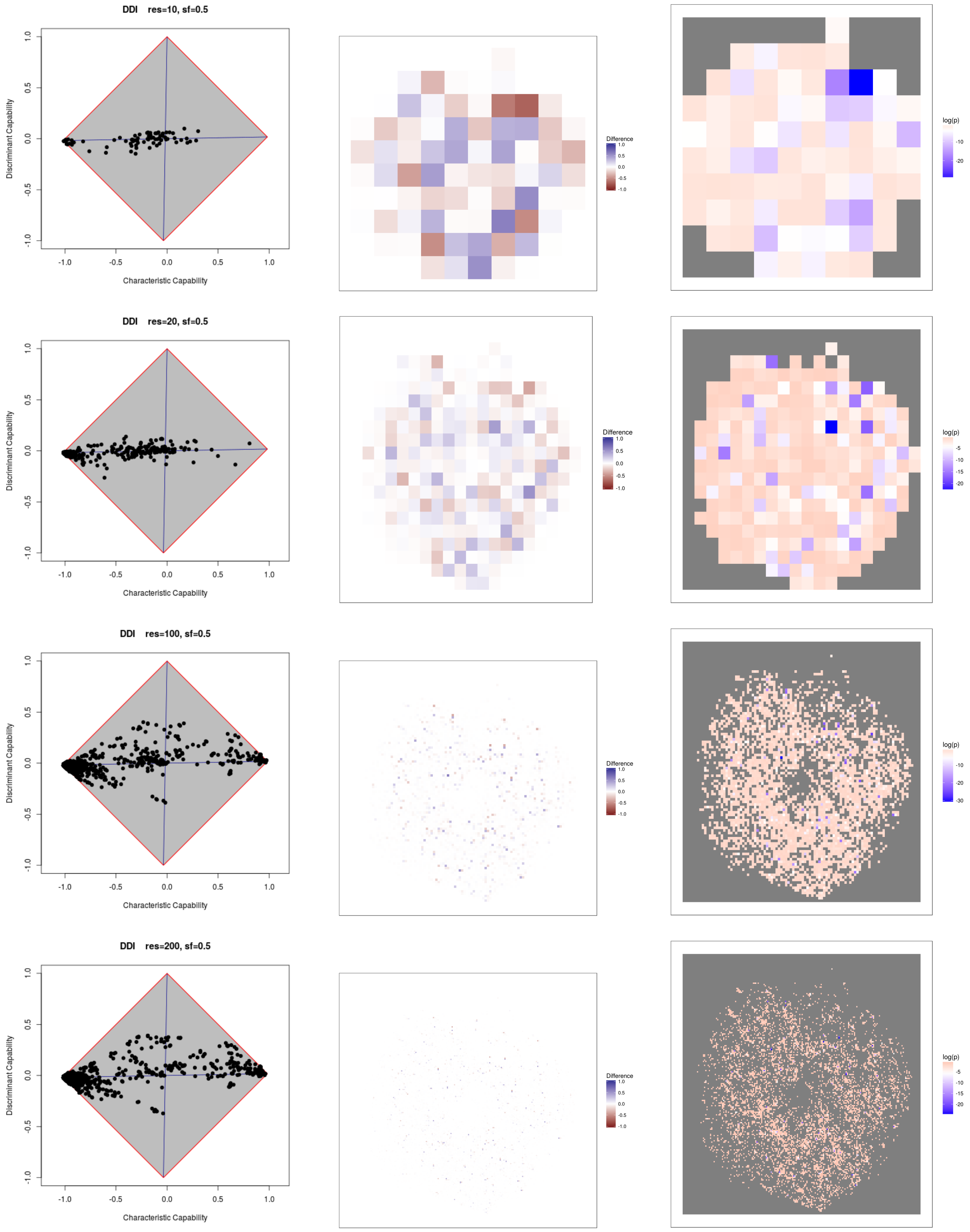


Figure 16: DDI, sf=0.5

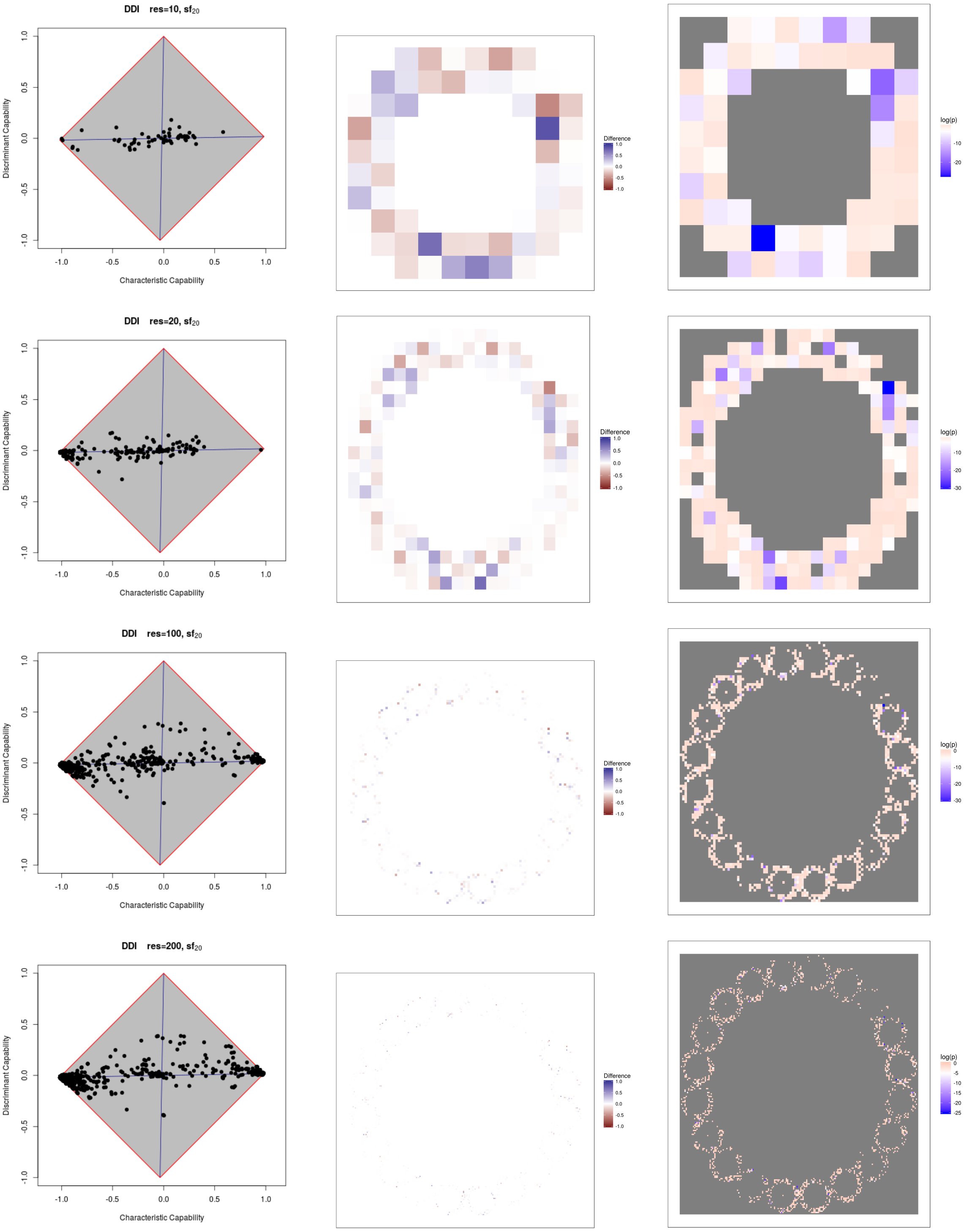


Figure 17: DDI, $sf_{20}$

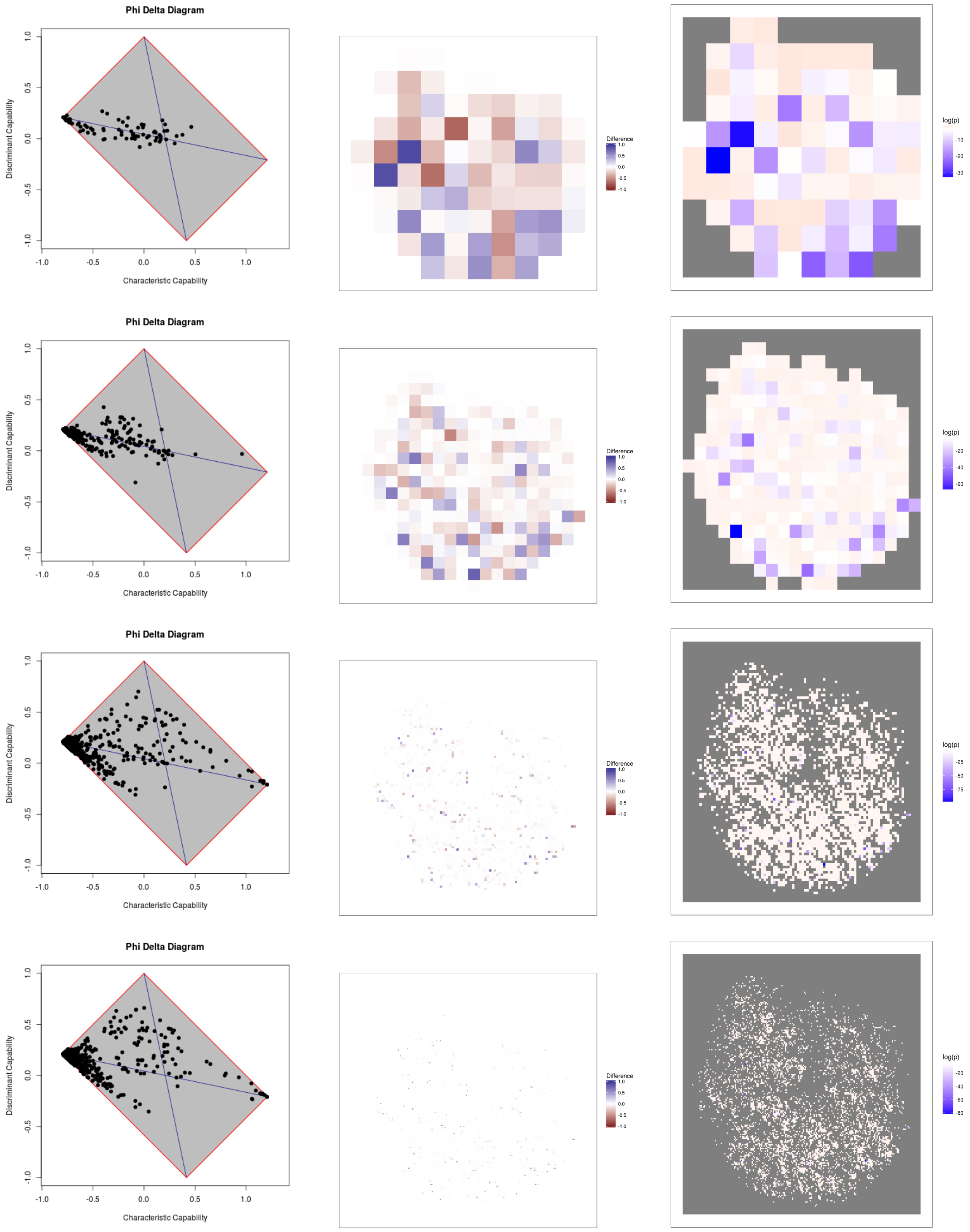


Figure 18: APV, sf=0.5

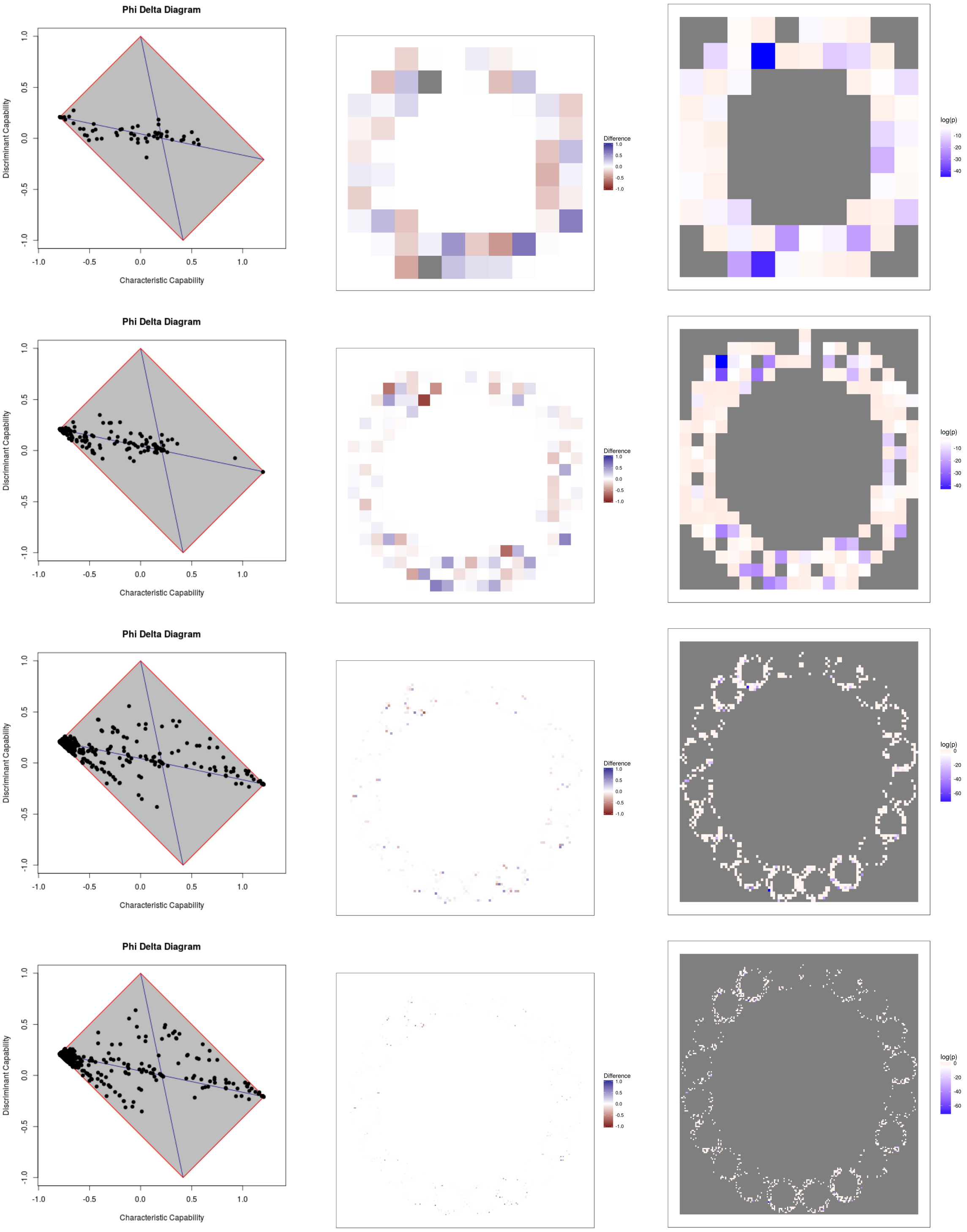


Figure 19: APV, $sf_{20}$

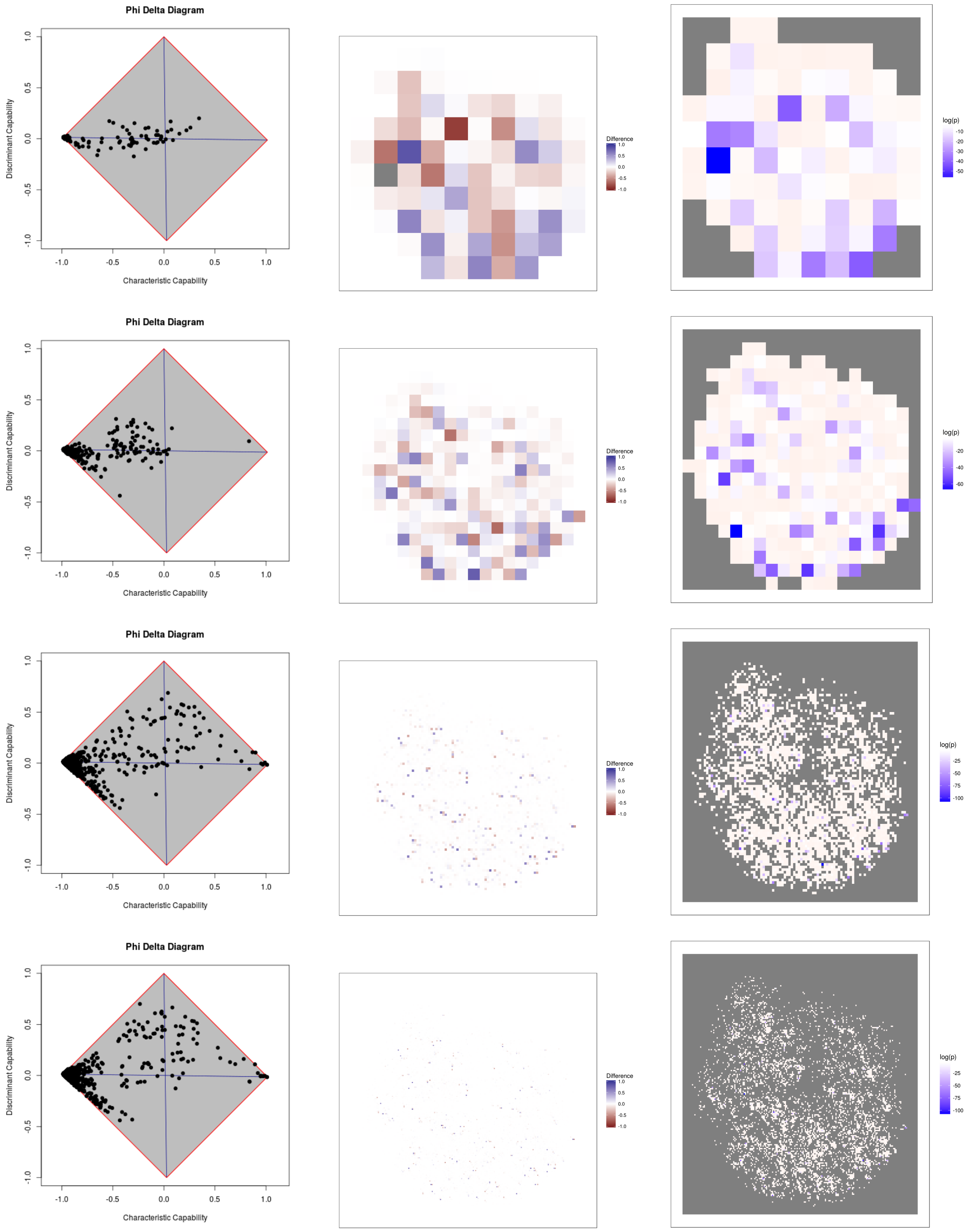


Figure 20: IDV, sf=0.5

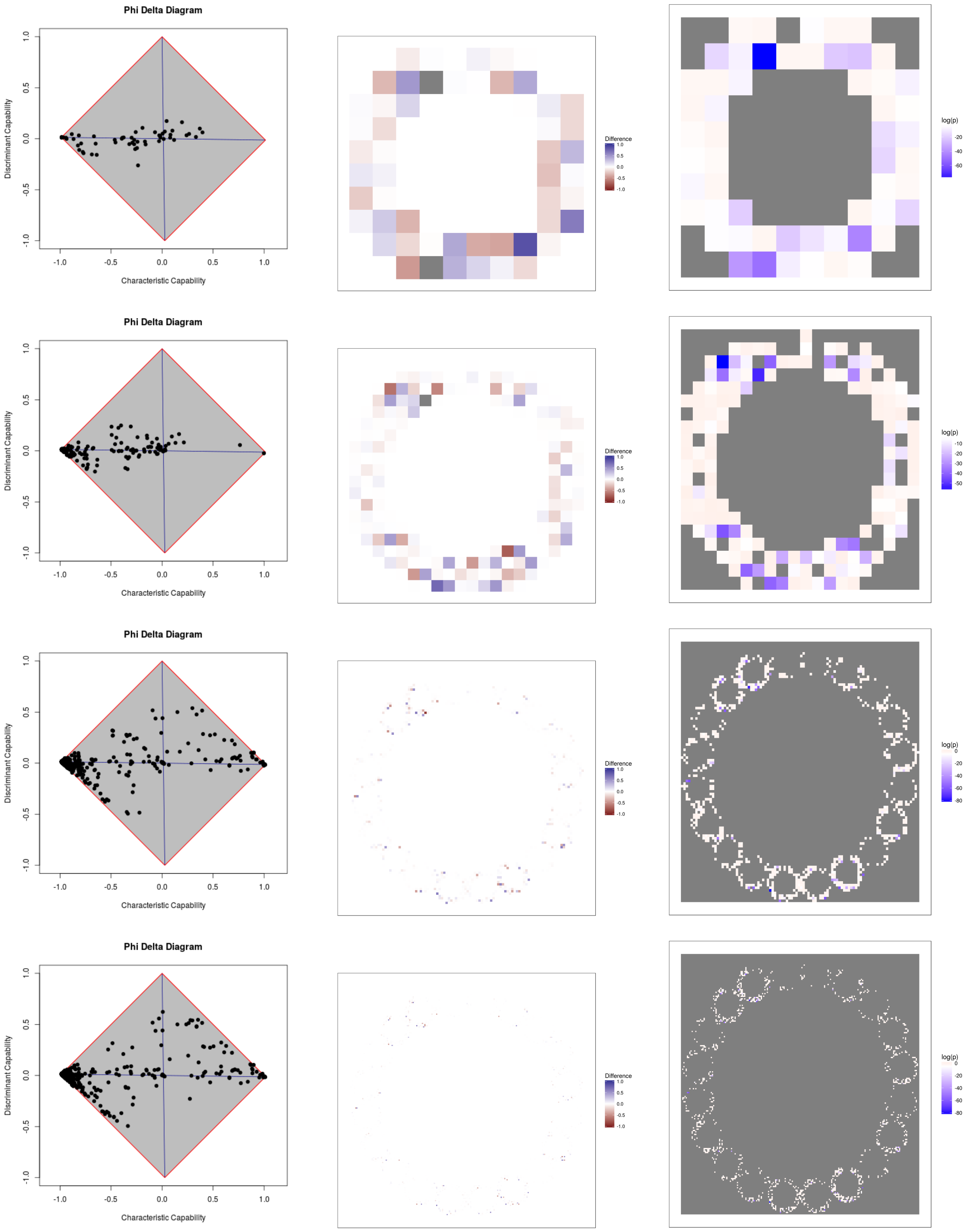


Figure 21: IDV, $sf_{20}$

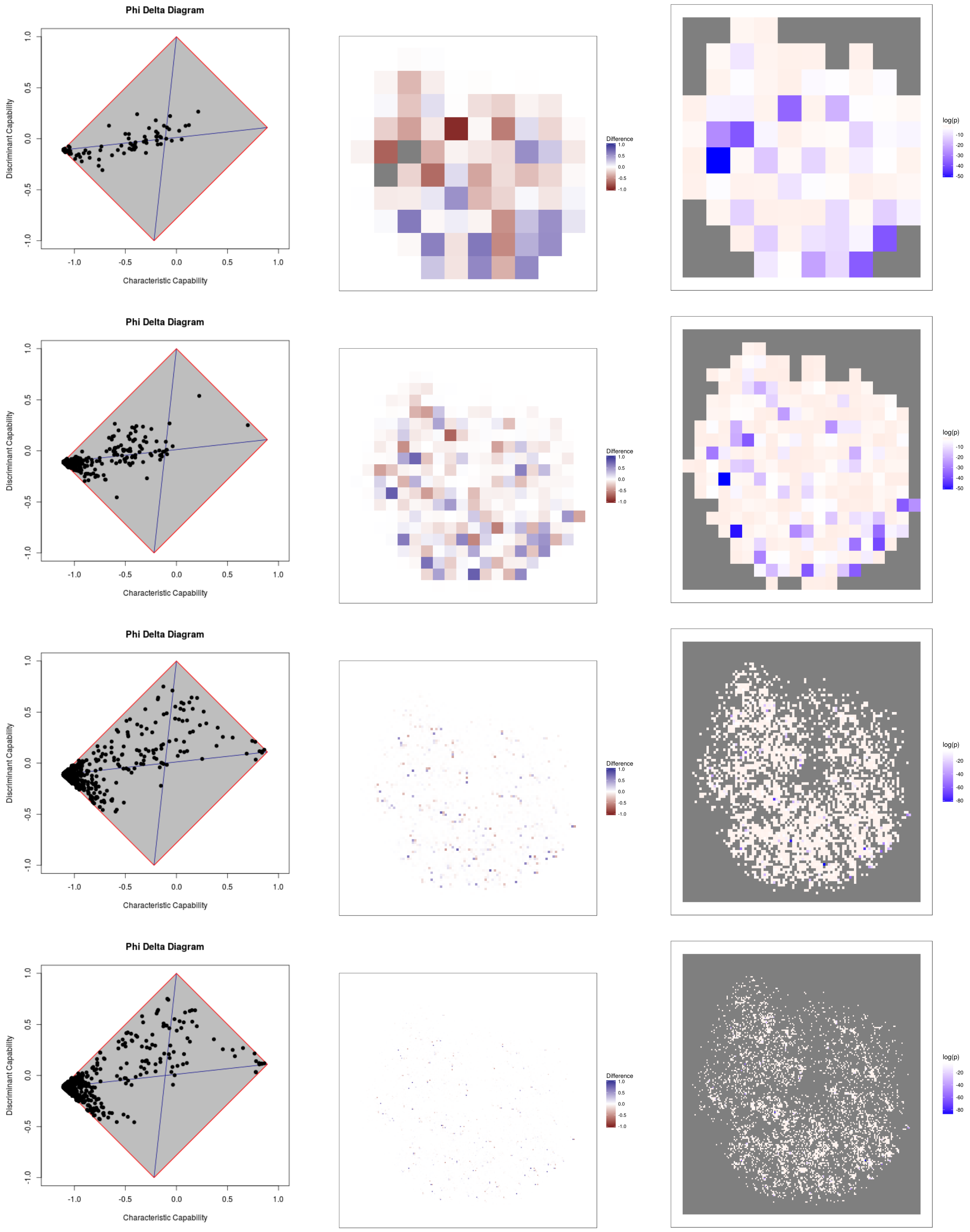


Figure 22: LPV, sf=0.5

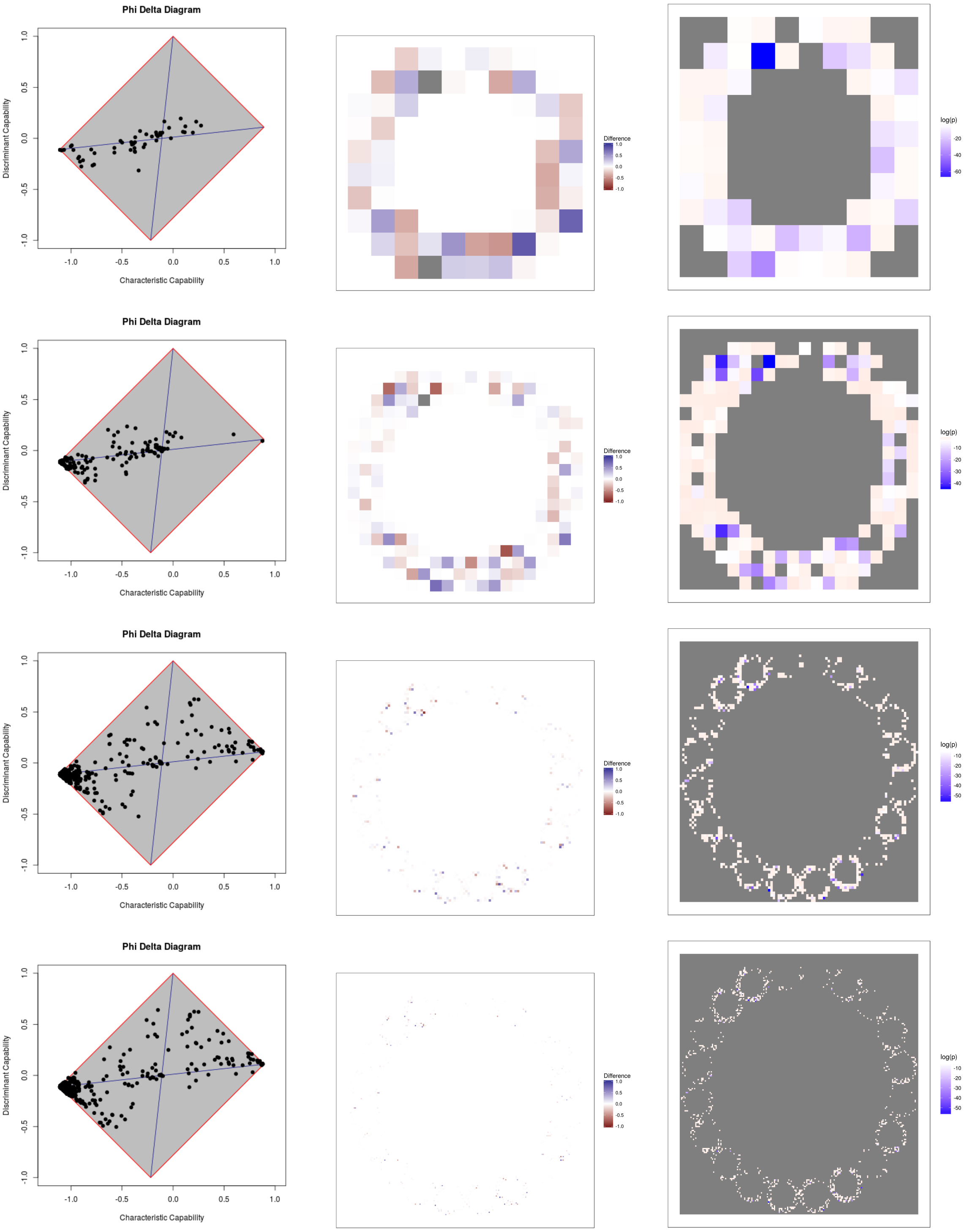


Figure 23: LPV, $sf_{20}$

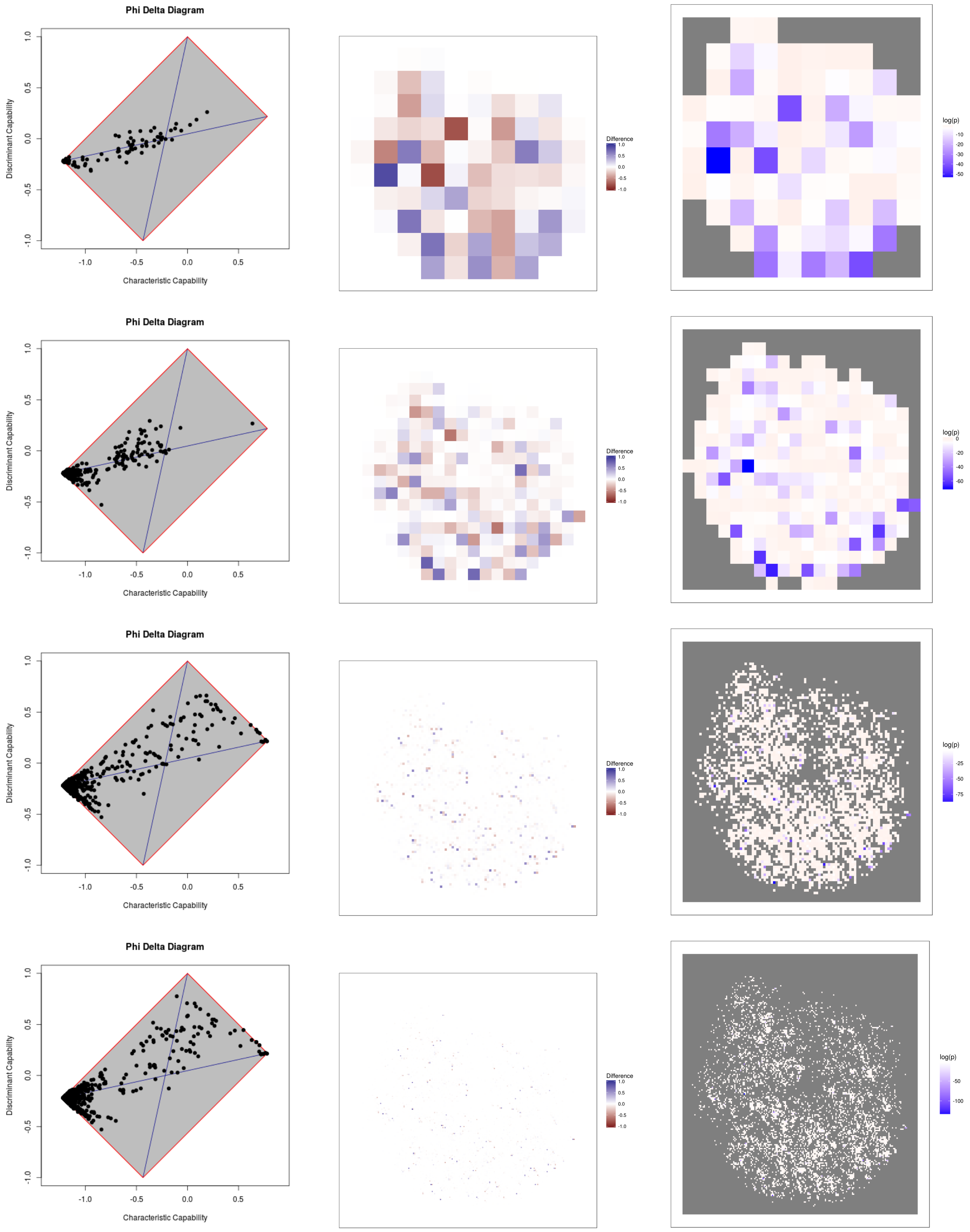


Figure 24: NFV, sf=0.5

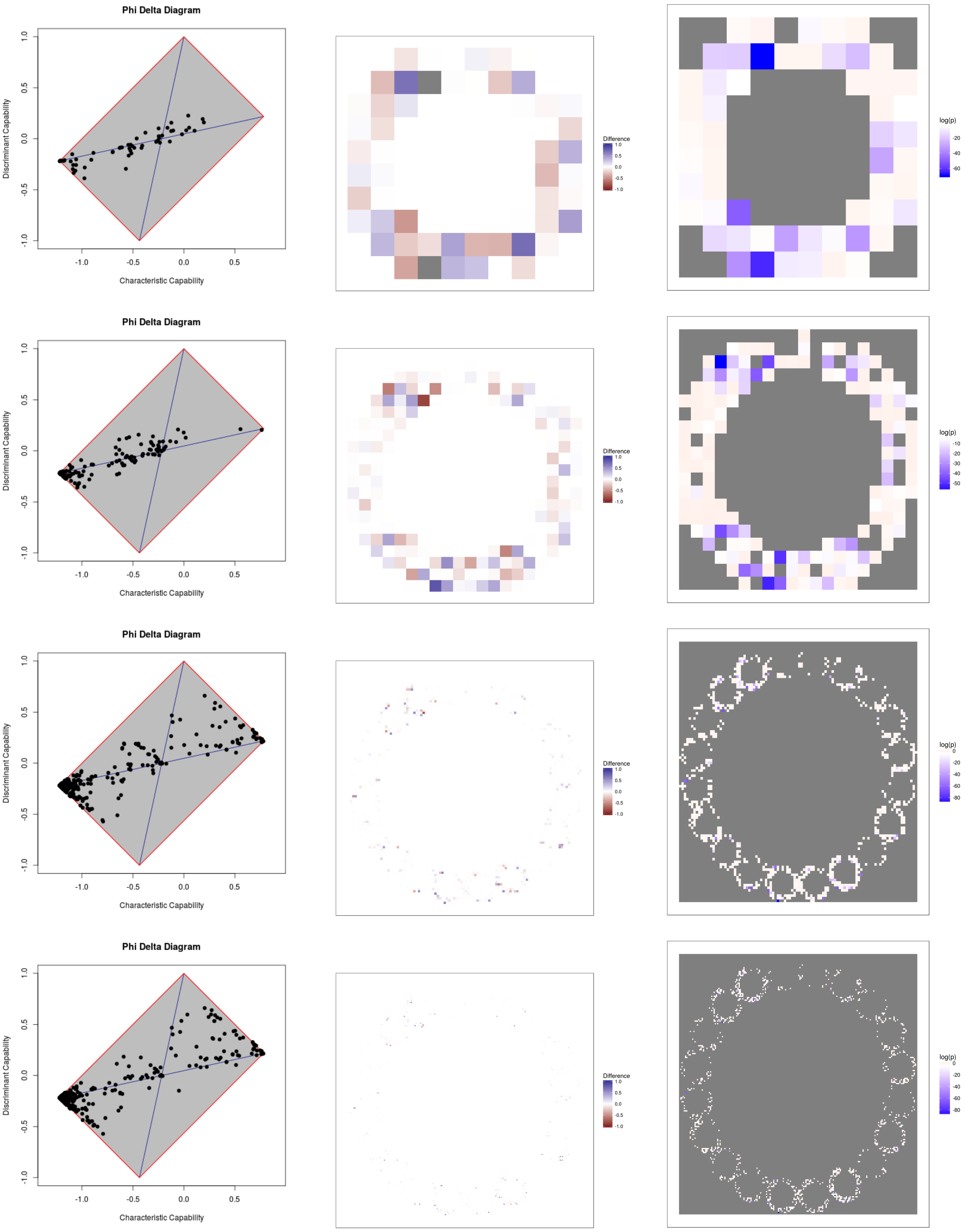


Figure 25: NFV, $sf_{20}$

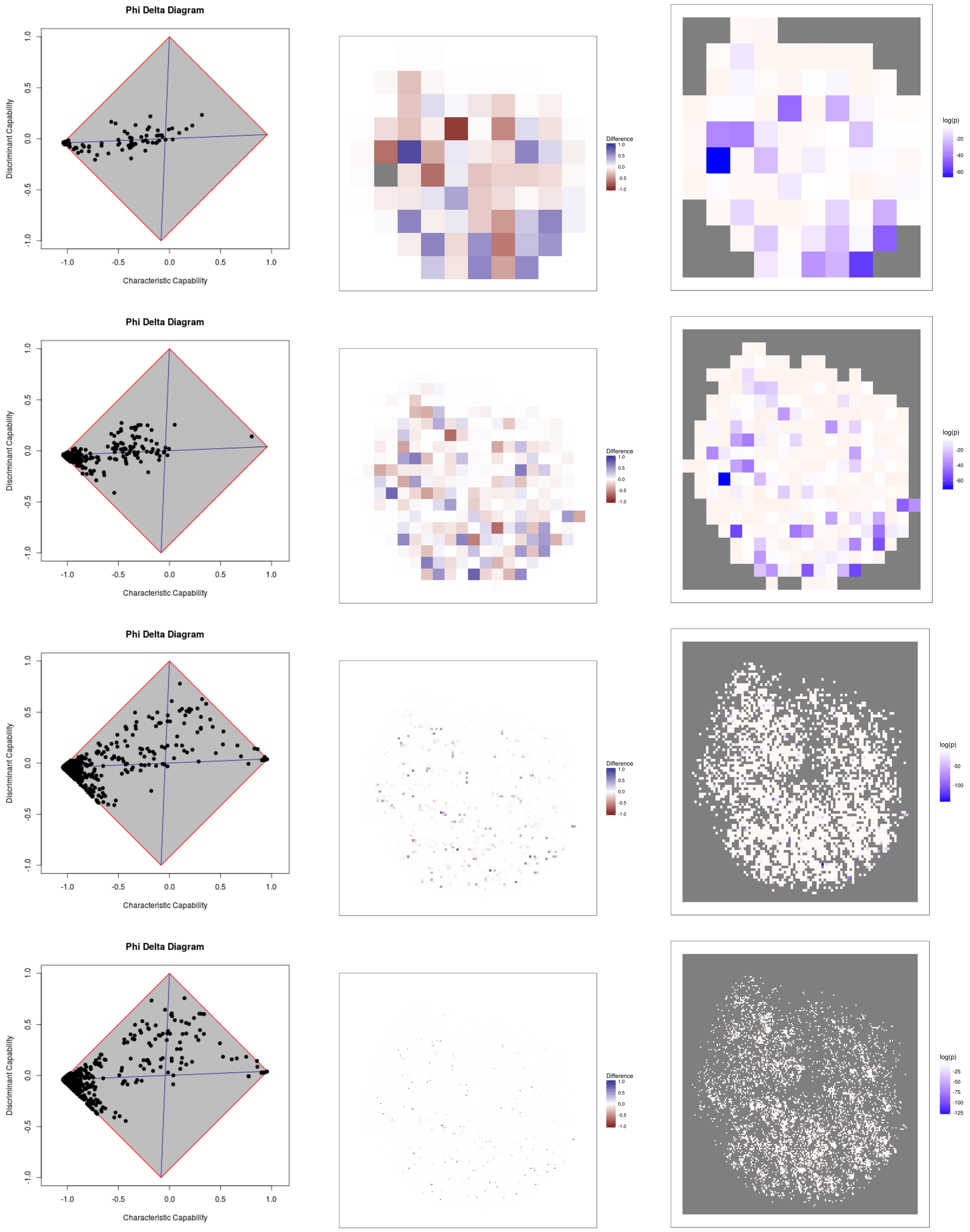


Figure 26: RTV, sf=0.5

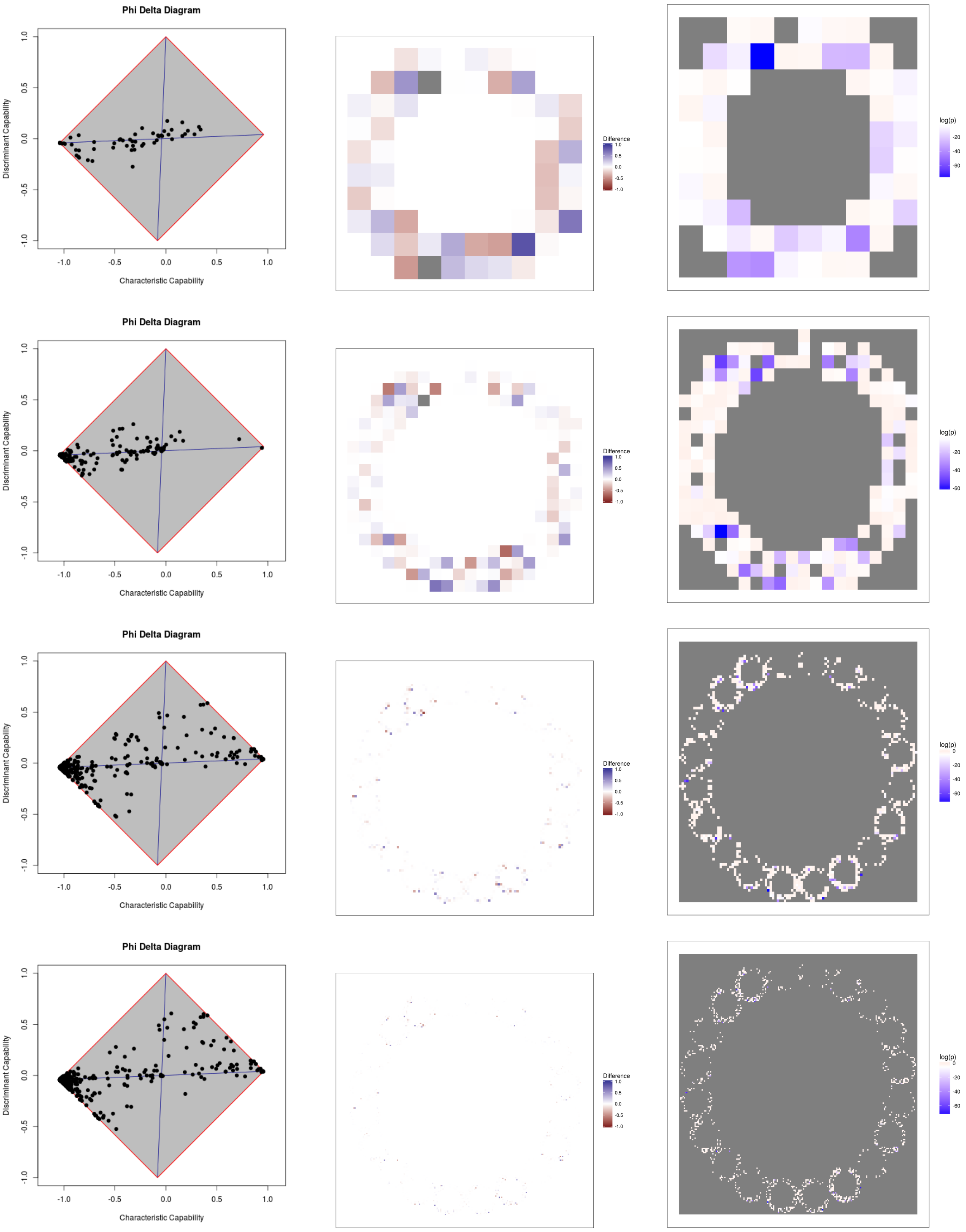


Figure 27: RTV, $sf_{20}$

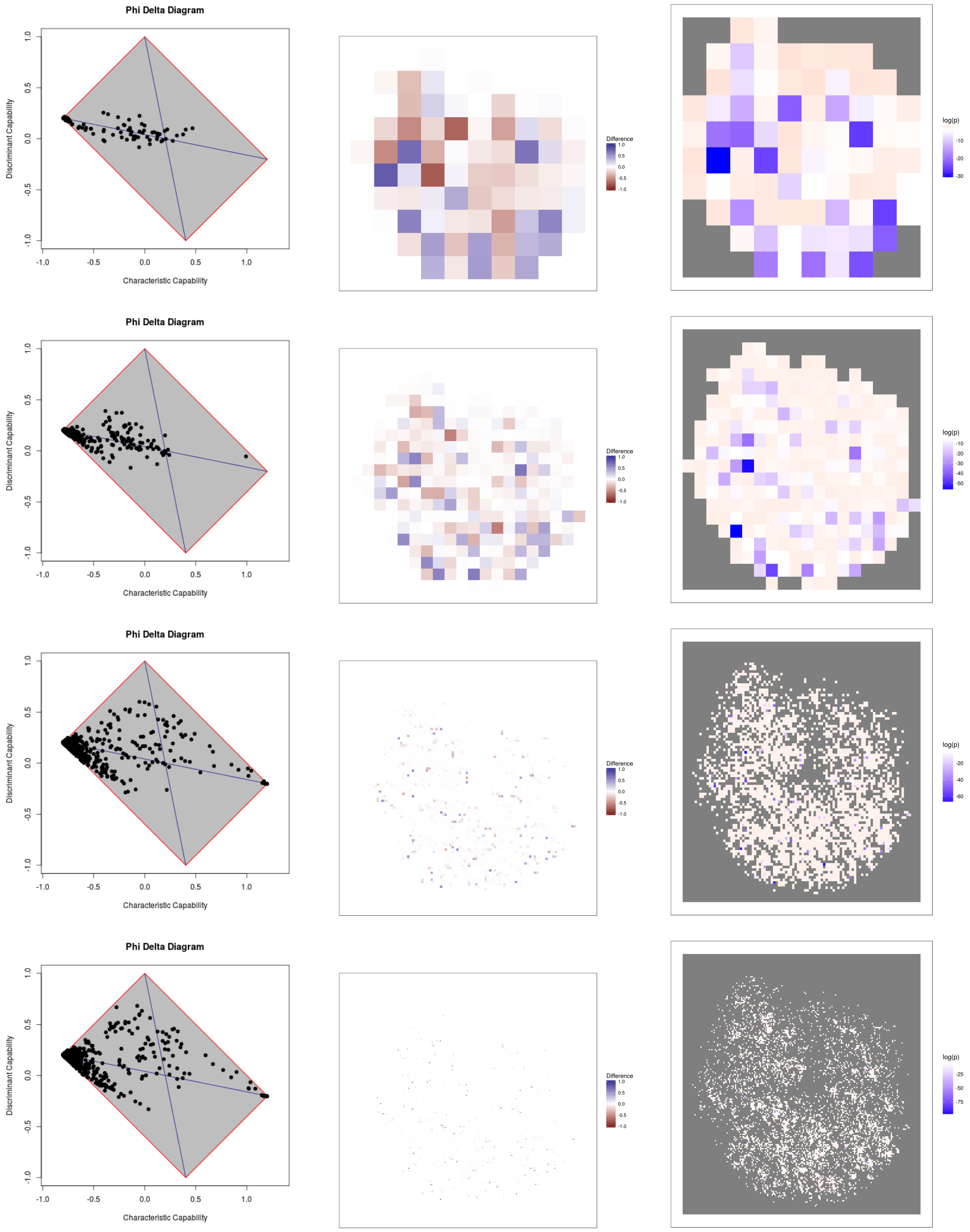


Figure 28: SQV, sf=0.5

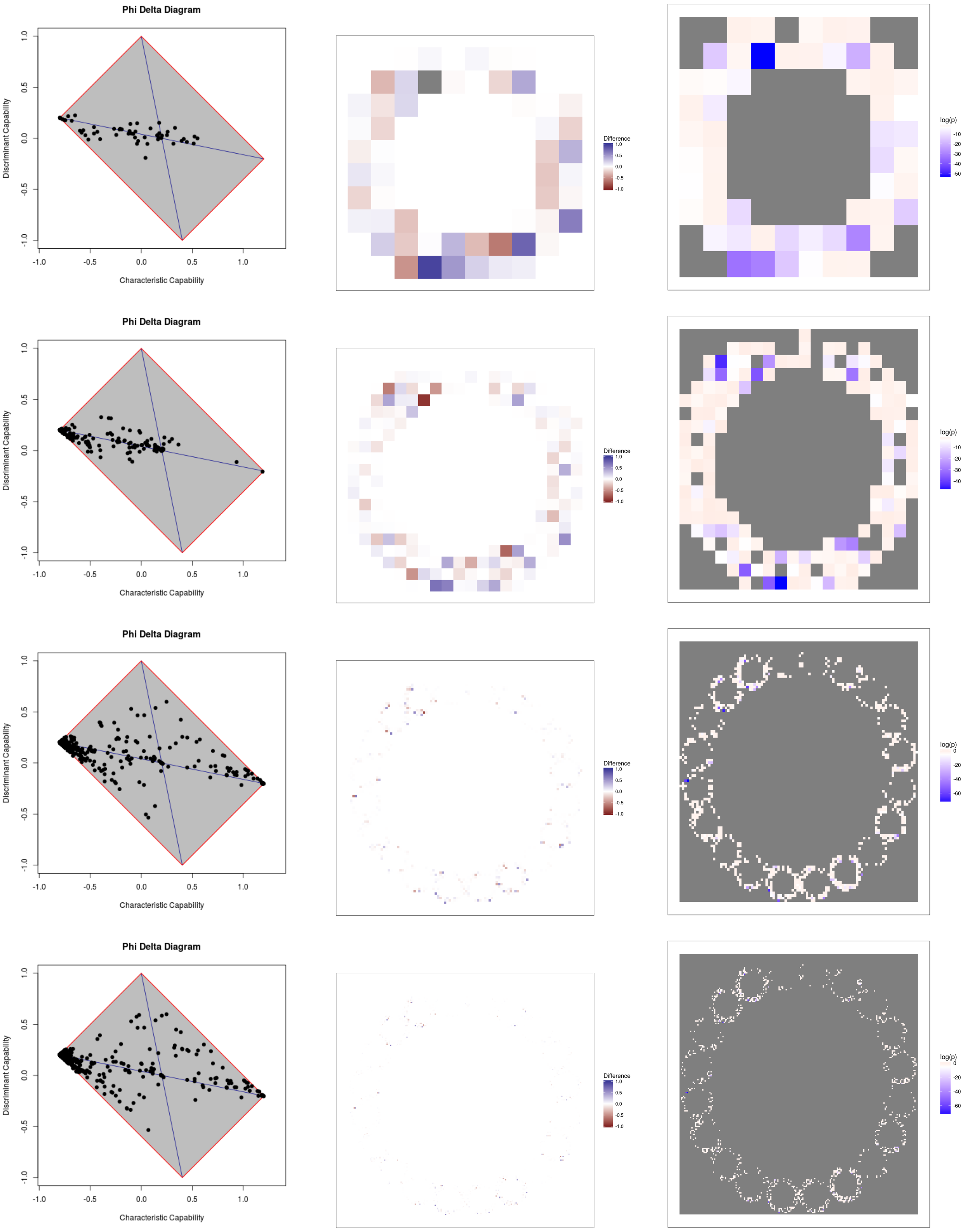


Figure 29: SQV, $sf_{20}$
